# Employing neutron-encoded monoUbs to study E2/E3 ligase activity and selectivity for assembling Ub chains

**DOI:** 10.1101/2025.05.22.655449

**Authors:** Bianca D. M. van Tol, Bjorn R. van Doodewaerd, Guinevere S. M. Lageveen-Kammeijer, Bas C. Jansen, Joanna Liwocha, Rishov Mukhopadhyay, Manfred Wuhrer, Gerbrand J. van der Heden van Noort, Brenda A. Schulman, Paul P. Geurink

## Abstract

While protein ubiquitination is an extensively studied post-translational modification, many aspects of this process remain unclear. Ubiquitin conjugation involves the action of three different types of enzymes working in concert to install ubiquitin onto substrate proteins. Despite efforts, an *in vitro* mid/high-throughput screen to quickly determine which enzymes work together to build ubiquitin chains and directly analyze the type(s) of chains formed does not exist. In this study, we developed a new multiplexed mass spectrometry-based E1-E2-E3 assay that enables the analysis of whether E2/E3 pairs work together to form ubiquitin chains and concomitantly reports on the nature of the formed ubiquitin chain type(s). The assay employs synthetic modified neutron-encoded monoUb substrates with a distinct molecular weight, enabling the simultaneous analysis of these substrates. Overall, various E2-E3 pairs were screened for their ability to build Ub chains, which furnished a three-dimensional overview of linkage selectivity over time and enzyme concentration.

**Graphical abstract:** 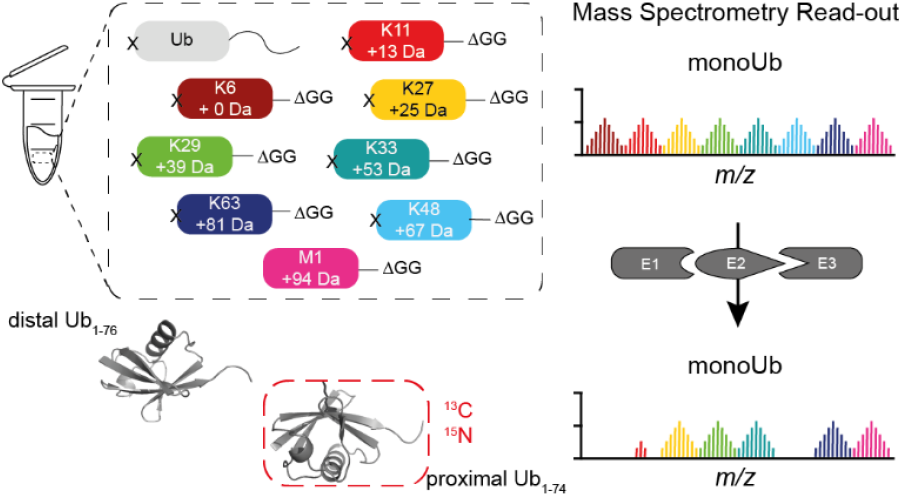

## Introduction

Ubiquitin (Ub)^1^ is a 76 amino acid protein that acts as a post-translational modifier, playing a key role in numerous cellular processes, including protein degradation, NF-κB signalling, DNA damage response, immunity, and inflammatory responses.^2,3^ Ubiquitination, the attachment of Ub to protein substrates, is a process that starts with the activation and adenylation of the C-terminus of Ub under the consumption of ATP by a ubiquitin-activating enzyme, E1, followed by the formation of a thioester-linked E1∼Ub complex.^4^ Ub is then transferred via a trans-thiolation onto the active site cysteine of a ubiquitin-conjugating enzyme (E2).^5^ Finally, with the help of a ubiquitin ligase (E3), Ub is attached via its C-terminus to a lysine residue side chain or N-terminal amino group of a substrate protein, forming a monoubiquitinated substrate.^6^ This process can be repeated on different lysines of the same substrate, resulting in multi-monoubiquitination, or on one of the seven lysines or N-terminus of Ub itself, to assemble polymeric Ub chains via any of the eight possible sites (Lys6, Lys11, Lys27, Lys29, Lys33, Lys48, Lys63 or Met1-linked). These polyUb chains can be homotypic when Ub is conjugated to the same site in each consecutive Ub. Mixed and branched chains can also arise when different positions are used during chain elongation. This whole repertoire of possible polyUb linkage types is often referred to as the Ub code.^2^ The human genome encodes two E1 enzymes^4^, around 40 E2 enzymes^5^ and 600 to 1000 E3 ligases^7,8^. The E3 ligases can be divided into three different families: Homologous to E6-AP C terminus (HECT), really interesting new gene (RING), and RING-between-RING (RBR) ligases.^9,10^ Each E3 ligase family uses a slightly different mechanism to support the E2 and the ubiquitination process. The E2 and E3 enzymes and their plethora of combinations determine substrate selection, the ubiquitination site (which lysine position(s) on the substrate), and the chain type and length, making the ubiquitination pathway a complex system.

With so many proteins involved, the Ub system is often dysregulated, leading to different diseases like cancer, inflammation, and neurodegenerative diseases.^11^ As each different (poly)Ub modification evokes different cellular responses, including protein degradation initiation, (de-)activation, localization, and changing interactions,^2,3^ it is important to elucidate which enzymes are responsible for the construction of Ub chain types shown that are involved in different disease pathogenesis, thereby creating opportunities for better targeted therapies. Therefore, one of the key challenges in the Ub field has been to determine which E2 and E3 enzymes work in concert and what type(s) of Ub chains they construct.

So far, various techniques have been used to address these questions (**Fig. 1a**). For example, SDS-PAGE analysis was used to show ligase-mediated chain assembly and/or autoubiquitination of ligases.^12,13^ However, the Ub chain type can only be determined by running single-lysine Ub mutant experiments side by side, or indirectly, by performing DUB restriction analysis (UbiCrest) on the products.^14^ While mass spectrometry has been pivotal in determining ubiquitination sites and chain topology,^15–18^ the ubiquitinated substrates need to be digested with trypsin to generate peptides that bear the 114 Da GlyGly modification on the ubiquitination site. Antibodies that detect Gly-Gly modifications on lysine residues^19^ or tandem ubiquitin-binding entities (TUBEs)^20^ were used for the enrichment of diGly-modified peptides or ubiquitinated substrates, which improved the amount of detected ubiquitination sites. Methods like absolute quantification (AQUA), which utilize diGly-modified ubiquitin-derived peptides, were used to quantify the Ub chain topology present in cell lysates and to determine which type(s) of Ub chains are built by a specific combination of ligase enzymes *in vitro*.^21,22^ Mass spectrometry techniques were also used to assess E2 and E3 ligases for their ubiquitin transfer activity, to identify active E2/E3 pairs, and to screen for E2/E3 inhibitors. Although these methods have significantly advanced knowledge on monoUb modifications, the chain types formed during the assay were not addressed.^23^ Furthermore, the Ub-clipping technique was designed to determine polyUb chain length and branching, but this required AQUA MS/MS to address the linkage type.^21^ Although all of these techniques provided valuable new insights into the assembly of Ub chains in cells and *in vitro*, they either do not allow for the determination of the Ub chain type or they require extensive and laborious sample preparation (digestion and enrichment).

**Figure 1.**
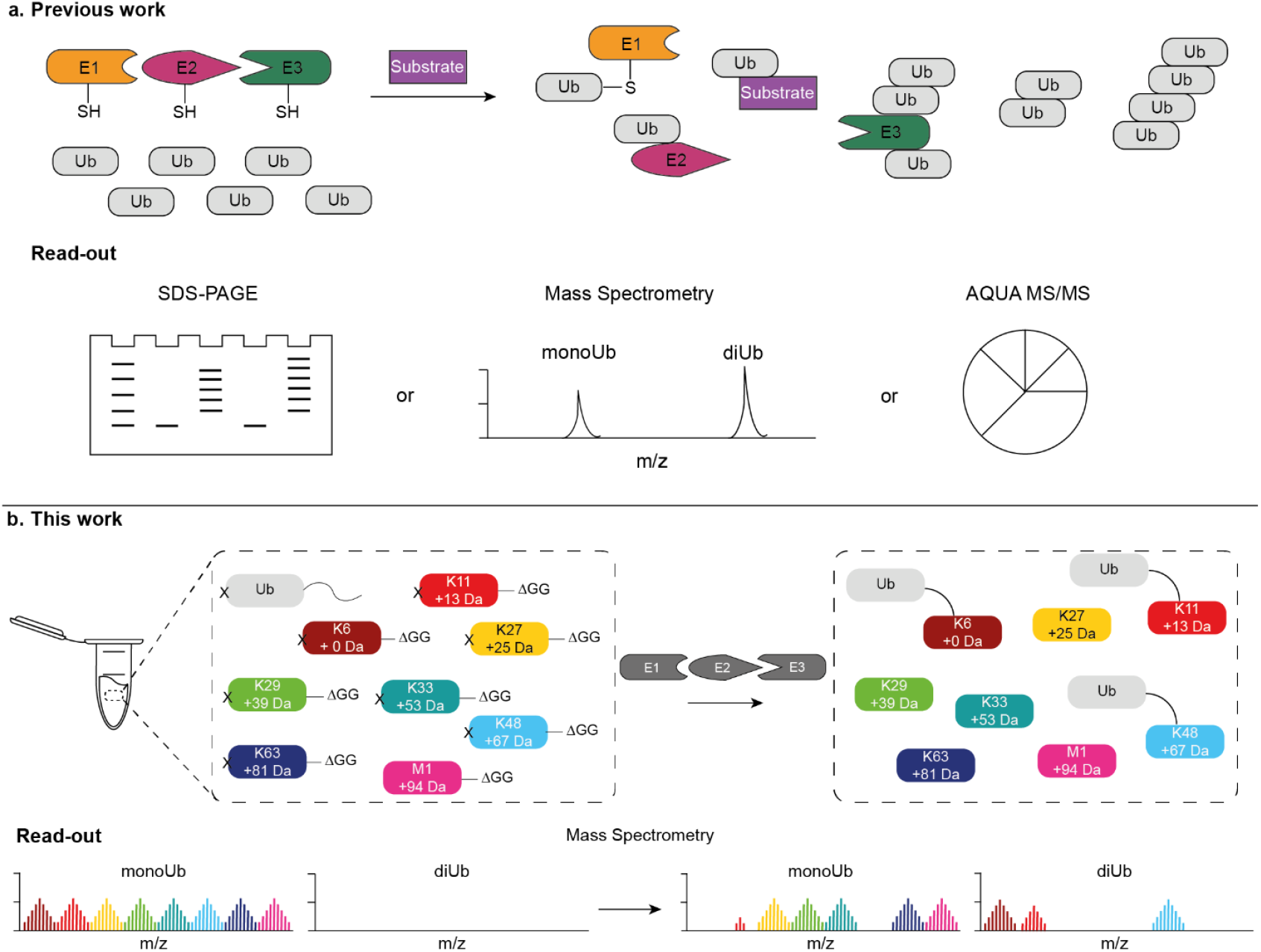
The principle of the neutron-encoded monoUb assembly assay. **a**, Schematic outline of classical ubiquitination assays where an E1, E2 and E3 enzyme are incubated with wild-type Ub (and a substrate). The products, free polyUb chains, ubiquitinated substrates, auto-ubiquitinated enzymes and enzyme∼Ub thioesters, are typically analysed by SDS-PAGE, mass spectrometry or AQUA MS/MS. **b**, Schematic outline of the designed method where an E1, E2 and E3 enzyme are incubated with a mixture containing synthetic modified neutron-encoded monoUb substrates. Subsequent MS analysis allows for the quantification of any of the eight possible produced diUb chains. Upon assembly, monoUbs which can form a specific (isopeptide)-linkage will disappear and the corresponding diUb signal will appear in the MS spectrum.

We recently developed a multiplexed mass spectrometry assay to analyse and quantify the deubiquitinase (DUB)-mediated consumption of all native diUb chains in competition with each other.^48^ The assay is fast, requires low amounts of substrates and provided a comprehensive overview of diUb chain processing, revealing new insights into the selectivity and chain processing order of different DUBs. Building on these results, we now adapted this assay to identify cooperating E2-E3 pairs to build (di-)Ub chains, and quantify the ratio of the formed diUb chain types, in a single read-out (**Fig. 1b**). We employed neutron-encoded single-lysine monoUb mutants, synthesized by solid-phase peptide synthesis (SPPS) (**Fig. 2a,b**), as substrates that can be decorated with a donor Ub. The donor Ub can be activated by an E1 enzyme at the expense of ATP and, via an E2 (and E3) enzyme, conjugated to the single-lysine monoUb mutants. Our assay measures and quantifies the consumption of different monoUbs and the formation of diUbs, distinguishing them in a single MS read-out by their different masses, resulting from the incorporation of neutron-encoded amino acids. We demonstrate that well-known E2 enzymes and E2/E3 pairs build their preferred diUb chains and show that, unlike DUBs,^24,25^ E2/E3 ligases do not show a concentration-dependent selectivity in the screened concentration range. We also explored an E2 panel to check whether these conjugating enzymes can independently form chains, and to identify the types of chains produced. Additionally, we used the assay to screen which E2 enzymes cooperate with two different E3 RING ligases by analysing whether diUb chains are formed and which type of chains are formed. The assay’s rapid read-out and multiplexed substrates streamline the process of determining enzymatic interactions and chain types produced. Notably, this assay requires low amounts of substrate, and the read-out can be quantified. Overall, this new assay unlocks valuable opportunities for studying enzymatic activity and selectivity in the Ub conjugation cascade.

**Figure 2.**
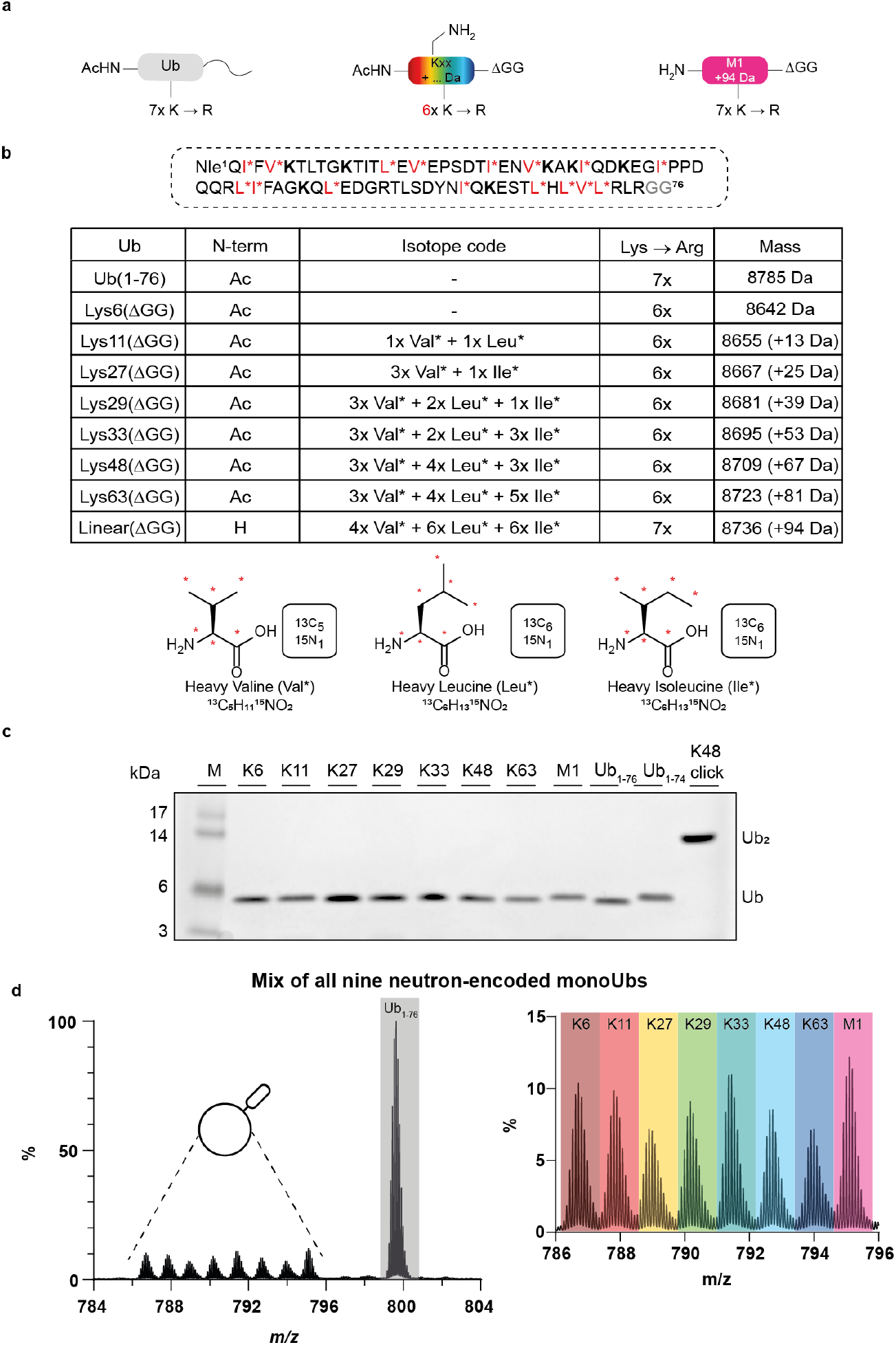
Design and synthesis of all monoUb substrates. **a**, Schematic overview of the modifications of the different monoUb substrates. Left: donor Ub with an acetylated N-terminus and seven K→R mutations. Middle: all monoUb acceptors which can form a single iso-peptide linkage with six K→R mutations, an acetylated N-terminus and a ΔGG modification. Right: the acceptor monoUb as precursor for linear chains, with a free N-terminus, seven K→R mutations and a ΔGG modification. **b**, Labelling scheme of the nine monoUb substrates. Potentially heavy-isotope labelled amino acid positions in the Ub sequence are red and marked with an asterisk. Nle = Norleucine. Positions for linkage-dependent K→R replacements are shown in bold. The table shows the number of introduced neutron-encoded amino acids and the introduced mass difference for each linkage. The isotope-labelled amino acids used are shown with the ^13^C and ^15^N atoms marked with red asterisks. **c**, Coomassie-stained SDS-PAGE analysis of all nine neutron-encoded monoUbs and used internal standards Ub1-74 and Lys48 click diUb. **d**, MS spectrum of a mixture containing equimolar amounts of all acceptor Ubs and an eight-fold excess of donor Ub, magnified at charge state z=11^+^.

## Results

### Design and synthesis of neutron-encoded monoUb molecules

For the designed MS-based assay (**Fig. 1b**), we required nine different monoUb substrate molecules. One acts as the donor ubiquitin, which is activated by the E1 enzyme and transferred through the E2 and E3 enzymes onto one of the eight acceptor substrates. The resulting products can then be analysed by mass spectrometry. To ensure that the donor Ub functions only as a donor, all of its lysine residues were mutated to arginines, and the N-terminus was blocked with an acetyl group to prevent the conjugation of a Ub onto this protein (**Fig. 2a, left**). Eight acceptor monoUbs had their C-termini truncated by two glycine residues (ΔGG) to prevent activation and loading onto the E1 enzyme. Six of the seven lysines were mutated to arginines, and the N-terminus was blocked by acetylation. (**Fig. 2a, middle**). For the monoUb serving as the acceptor for linear diUb formation, all lysines were mutated to arginines but the N-terminus was kept free (**Fig. 2a, right**). Furthermore, neutron-encoded amino acids were incorporated to introduce mass differences of 12, 13 or 14 Da among the molecules, facilitating a proper isotopic pattern separation in the MS spectrum.^48^ Valine, Leucine and Isoleucine residues were replaced by their fully ^13^C and ^15^N labelled isotopologues, introducing a mass difference of 6 or 7 Da per substitution. Details of the substitutions are shown in **Fig. 2b** and **Supplementary Table 1**. The mass difference was specifically introduced in the acceptor monoUbs to link a mass difference to a specific type of ubiquitin chain that can be formed. The monoUb substrates were synthesized using linear solid-phase peptide synthesis (SPPS), allowing for amino acid modifications (**Supplementary Scheme 1**). After linear SPPS, the proteins were liberated from the resin under acidic conditions and purified by RP-HPLC and size exclusion. Their purity was confirmed by SDS-PAGE (**Fig. 2c** and **Supplementary Figure 1**) and LC-MS (**Supplementary Data**) analysis.

### All monoUb substrates can be measured simultaneously, and modified substrates can be processed by ligase enzymes

To assess the possibility of analysing all nine monoUbs simultaneously, and whether they could be used in a multiplexed MS assay, the acceptor molecules were mixed in an equimolar ratio with one equivalent of donor molecule for each acceptor, to make it theoretically possible to convert all monoUbs into diUbs. This resulting mixture was analysed by HPLC-MS, showing a proper baseline separation of all monoUb molecules (**Fig. 2d** and **Supplementary Figure 2**). The eight-fold excess of donor Ub was clearly detected, while the lower-abundant monoUbs gave a well-detectable signal. We confirmed that these modified substrates were still recognized and correctly processed by the ubiquitination enzymes. The donor Ub could still be loaded onto the E1 and E2(-E3) ligase, as confirmed by the formation of the thioester-linked E1∼Ub and E2(-E3)∼Ub conjugates that became apparent upon SDS-PAGE analysis under non-reducing conditions (**Fig. 3**, lane 3 in both gels). Under the same conditions, the acceptor monoUbs were not loaded onto the E1 and E2(-E3) enzymes (**Fig. 3**, gel left, lane 4). Furthermore, a selective K48 Ub linkage forming E2/E3 fusion ligase of Ube2G2 and gf78^26^ was used to check whether K48 diUbs could still be formed (**Fig. 3**, gel left, lane 5-8), and whether other chains, for example K6, were not formed when all other Lys positions are not available for ligation (**Fig. 3**, gel left lane 9-11). This fusion ligase was able to build diUb (band around 14 kDa), triUb (band around 21 kDa), and even polyUb chains (protein smear) when wild-type Ub was used as substrate (**Fig. 3**, gel right, lanes 4-7). Moreover, the formation of diUb also occurred when all eight different acceptor substrates and the donor Ub were present, as confirmed by the protein band around 14 kDa (**Fig. 3**, gel right, lanes 8-11). However, from this band on gel, it was not possible to determine which chain type(s) were formed, as that can only be determined using MS (*vide infra*). The activation and conjugation reactions are all ATP-dependent, as became clear from **Figure 3** (Gel left, lane 8 and Gel right, lane 7 and 11), since no product formation was observed in the absence of ATP.

**Figure 3.**
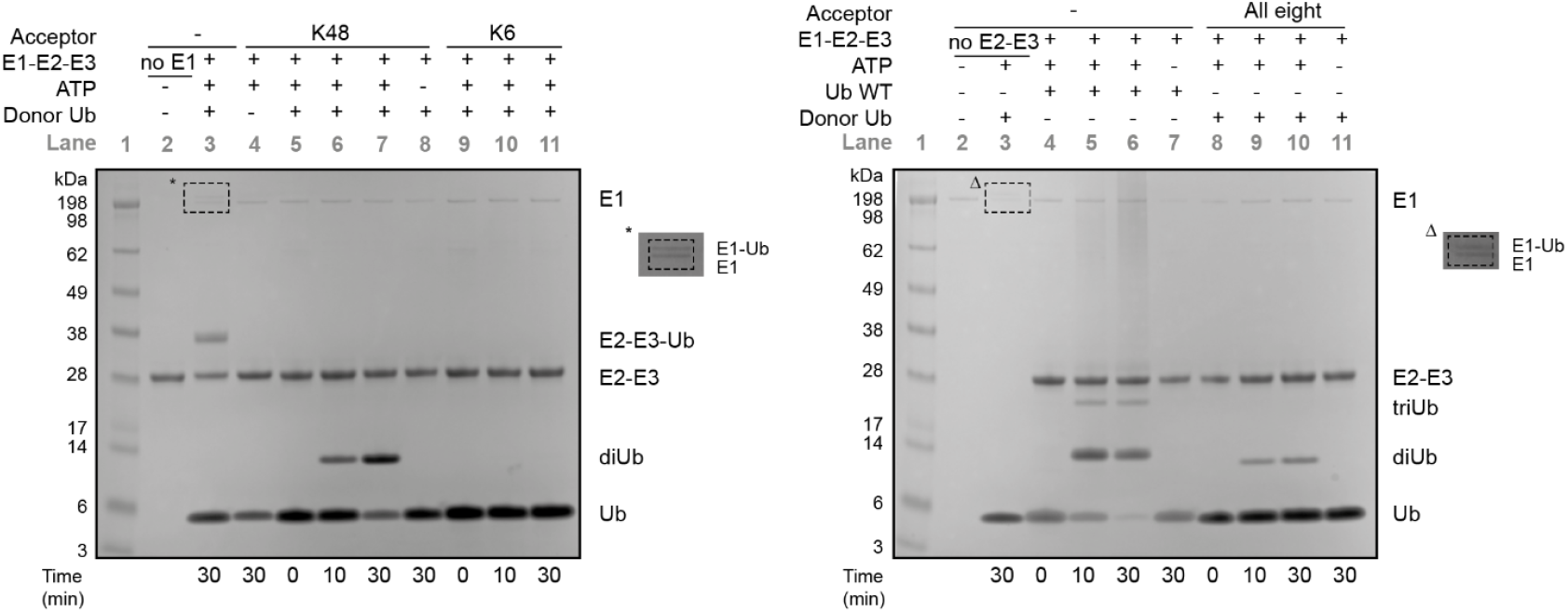
Proof of Principle Assay. Coomassie-stained SDS-PAGE analysis of ubiquitination assay. Ub WT, donor Ub, and different acceptor Ubs were incubated with UBE1, and/or an E2-E3 fusion enzyme of UBE2G2 and gf78. E1-Ub and E2-E3-Ub thio-ester formation was monitored under non-reducing conditions (left gel, lane 3 and right gel, lane 3), and diUb and polyUb formation were monitored under reducing conditions.

### Determining diUb linkage formation by linkage-specific assembly enzymes

After confirming that all monoUbs can be detected in a single mixture and that they are appropriately processed by ligase enzymes, we investigated whether we could make all different diUb molecules using reported specific enzymes or enzyme pairs.^12,22,27–30^ For this assay, a mixture containing all eight acceptor monoUbs (5 μM each) and eight equivalents of the donor monoUb (40 μM) was prepared, and subsequently mixed with a solution containing recombinantly expressed UBE1 (100 nM), UBE2x (2 μM, 6 μM or 10 μM) and E3 ligase (4 μM, 12 μM or 20 μM) at 37°C. The production of diUb was initiated by the addition of ATP. Samples were taken at various timepoints (at t = 0, 10, 30, 60, 180 minutes), quenched by adding the sample to an acidic solution containing the internal standards, and subsequently analysed by LC-MS. Ub_1-74_ and non-hydrolysable clicked Lys48 diUb were chosen as internal standards to control for chromatogram alignment. Quantification was performed over the whole charge state range of the proteins (z=10^+^ to z=25^+^ for diUb and z=5^+^ to z=13^+^ for monoUb) using a tailor-made version of the open-source software package LaCyTools.^31,32^

We started with the formation of Lys6, Lys11, Lys29, Lys33, Lys48, Lys63, and Met1-linked diUb chains to check whether we could make ‘all’ diUb chains (**Fig. 4a** and **Supplementary Figure 4**). An enzymatic *in vitro* assembly E2(-E3 pair) for Lys27-linked polyUb chains has not been described yet, so it was not possible to check the formation of this linkage. The combination of UBE2L3 and NleL effectively produced a mixture of Lys6-(∼66%) and Lys48-linked (∼34%) diUb chains already after 10 minutes. We opted to produce Lys11 linkages with the E2 construct UBE2S-IsoT, which has been shown to be more efficient in producing ubiquitin multimers than the native UBE2S enzyme.^29^ Unfortunately, no Lys11-linked or other diUbs were produced during the assay. To check whether the UBE2S-IsoT enzyme was indeed active, we analysed Ub chain formation using Ub WT and donor Ub and the K11 acceptor solely by SDS-PAGE analysis (**Supplementary Figure 6**). DiUb formation was observed for Ub WT and autoubiquitination of UBE2S-IsoT was observed for both Ub WT and the modified donor Ub, meaning that the enzyme is active and able to accept the modified donor Ub. Again, no diUb formation was observed with Lys11 acceptor Ub. To show that the production of Lys29-linked chains was possible, we used the combination of UBE2L3 and UBE3C, which produced a mixture of Lys-6, Lys-29, and Lys-48-linked ubiquitin chains (∼3%, ∼17%, ∼80%, respectively). UBE2L3 in combination with AREL1 was used to produce Lys33- and Lys48-linked diUb chains, which showed the formation of around 88% of Lys33-linked diUb. Although we already proved that it is possible to assemble Lys48-linked diUb chains with our assay using other enzyme combinations, we opted to show that we could solely produce this diUb chain type by using either UBE2G1, UBE2R1 or UBE2R2. These three E2 conjugating enzymes produced solely Lys48-linked diUb constructs, and the total amount of formed Lys48-linked diUb showed to be dependent on the assay time, E2 concentration, and the type of E2 enzyme (**Fig. 4a** and **Supplementary Figure 4**). Lys63-linked diUb chains were readily formed by the E2 pair UBE2N and UBE2V1. UBE2V1 is a ubiquitin E2 variant, which lacks the active-site Cys-residue but interacts with UBE2N to form Lys63-linked polyubiquitin chains specifically. Finally, Met1-linked or linear diUb chains could be produced by UBE2D3 in combination with HOIP. Although the production of linear diUb chains was slow, after 180 minutes, some linear diUb could be observed, and overnight reaction resulted in a substantial amount of linear diUb. These assays proof that despite the modifications made to the monoUb’s (e.g. K→R mutations, acetylation, ΔGG) they are still properly recognized by the E1-E2-E3 enzymes and turned into the expected diUb linkages starting from the mixture of donor monoUb and all eight acceptors present.

**Figure 4.**
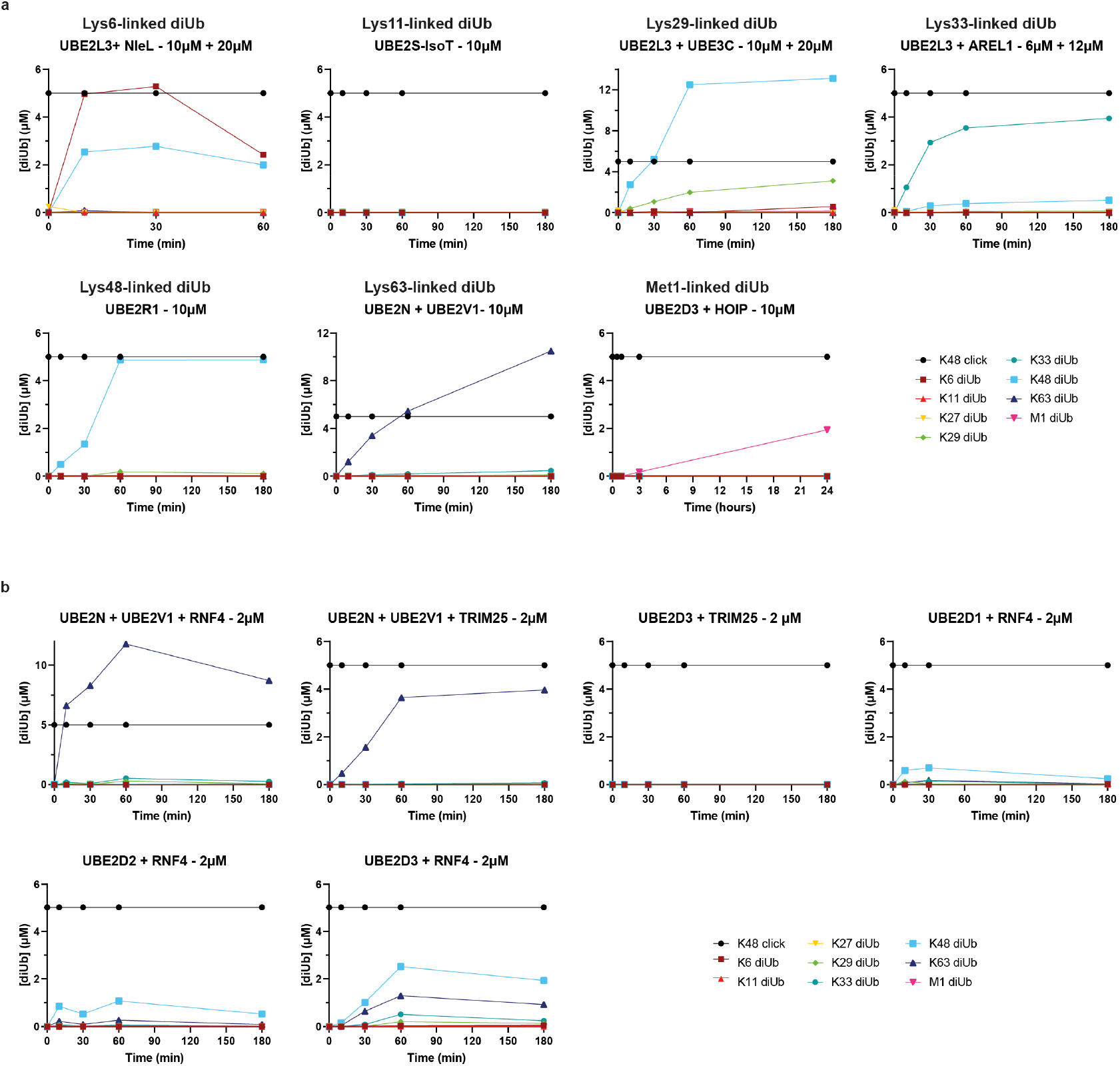
Assembly of differently linked diUb chains. **a**, Assembly of Lys6-, Lys11-, Lys29-, Lys33-, Lys48-, Lys63-, and Met1-linked diUb chains over time using different E2 enzymes or E2/E3 enzyme pairs. A mixture containing all eight acceptor monoUbs (5 μM each) and donor monoUb (40 μM) was added to a solution containing recombinantly expressed UBE1, UBE2x, and E3 ligase at 37°C. The production of diUb was initiated by the addition of ATP and analyzed by LC-MS. **b**, Screen of different E2/E3 enzyme pairs for their diUb chain assembly activity and selectivity.

E3 RING ligases can bring an E2 enzyme and a substrate together to speed up the ubiquitination process.^7^ To show that our assay can be used to assess this, we took the UBE2N + UBE2V1 combination and checked whether the formation of Lys63-diUb chains was accelerated by the addition of E3 ligases RNF4 or TRIM25. Indeed, with the addition of RNF4 to the enzymatic cascade, the same amount of Lys63 diUb was produced within 10 minutes at 2 µM of E2 enzyme, compared to 60 minutes at 10 µM in the absence of RNF4 (compare **Fig. 4b** with **Fig. 4a** – Lys63-linked diUb). The addition of TRIM25 increased the Lys63-linked diUb formation rate by around 1.3 times. TRIM25 has also been reported to bind to UBE2D3, so we tested whether UBE2D3 and TRIM25 cooperate to build diUb chains. The assay revealed that after 180 minutes at high enzyme concentration, no diUb chain formation was observed. However, the consumption of the donor Ub in the monoUb graph was clearly visible, likely arising from TRIM25 autoubiquitination (**Supplementary Figure 5**), which corroborates reported findings.^33,34^

### Determining *in vitro* chain formation by UBE2Dx

The four UBE2Dx enzymes (UBE2D1, UBE2D2, UBE2D3, and UBE2D4) are reported as a promiscuous class of E2 enzymes, which work together with many different E3 ligases^23^ and are interchangeable.^35,36^ However, UBE2Dx enzymes are hardly active on their own.^23^ Indeed, in our assay, we did not observe diUb chain formation for any of these UBE2Dx enzymes in the absence of an E3 after 24h incubation (**Supplementary Figure 6**). The addition of RING ligase RNF4 to the assay resulted in the production of a variety of diUb chains. For UBE2D1 combined with RNF4, mainly Lys48-linked chain formation was observed (**Fig. 4b**), and after 3 hours, Lys63-linked chains were observed at high enzyme concentrations (10 µM) (**Supplementary Figure 5**). UBE2D2 in combination with RNF4 also mainly built Lys48-linked chains, and the formation of Lys63-linked diUb chains was also observed, already at low enzyme concentration (**Fig. 4b**). At high enzyme concentrations, the production of Lys29- and Lys33-linked chains could be observed as well (**Supplementary Figure 5**). UBE2D3 in combination with RNF4 was more active than the other UBE2Dx members, and the formation of mainly Lys48-linked chains but also Lys63-, Lys29-, and Lys33-linked diUb chains were clearly detected in this assay within one hour. Based on these results, we hypothesize that the enzymes within the UBE2Dx class may possess a similar and somewhat promiscuous selectivity profile, but each with different chain-building efficiency.

### Determining *in vitro* chain formation of E2 enzymes

In addition to E3 ligases, E2 enzymes recently emerged as key mediators of ubiquitin chain assembly. Approximately 30 E2’s can conjugate Ub to an amino group themselves in the presence but also the absence of an E3 ligase,^38^ and they have an important role in the determination of the ubiquitin chain length and topology.^5,37^ In the absence of E3 ligases, typically high concentrations of E2 enzymes are needed, and transfer rates of ubiquitin onto a substrate are low.^37^ In 2018, De Cesare *et al*. screened the activity of E2 conjugating enzymes in the absence of E3 ligases by monitoring the consumption of the monoUb pool.^23^ However, the activity of E2 ligases can be split into Ub chain formation and auto-ubiquitination of the E1 and E2 enzymes. With their MALDI-TOF assay, the products were not analysed, and the fate of the monoUb was unknown. To analyse whether the monoUb is used for chain formation or auto-ubiquitination, we used our neutron-encoded monoUb set and tested 21 recombinantly expressed E2 conjugation enzymes (**Supplementary Table 2** and **3**) for their ability to build diUb chains at an enzyme concentration of 1 μM E2. After 1h and 24h, we analysed the amounts of monoUb and diUb present in the reaction mixture. (**Fig. 5a**). No significant diUb chain signals were observed after 1h (**Supplementary Figure 6**), but after 24h, we observed Lys48 diUb formation for UBE2G1, UBE2G2, UBE2K, UBE2R1, and UBE2R2, and Lys63 diUb formation for UBE2N-UBE2V1, which is in accordance with literature.^28,39,40^ MonoUb consumption data showed the consumption of Ub_1-76_ for UBE2A, UBE2G2, UBE2J1, UBE2K, UBE2Q1, UBE2Q2 and UBE2W. In the absence of diUb chain formation for UBE2A, UBE2J1, UBE2Q1, UBE2Q2 and UBE2W (**Fig. 5a**), these results are pointing in the direction of auto-ubiquitination of the E1 and E2 enzymes. However, since 8 equivalents of donor Ub are present in this experiment, and the measured amount of reaction mixture was optimized for the acceptor Ubs and the formed diUbs, the signal for the donor Ub analyte is relatively high and can therefore not be quantified accurately. Different amounts of reaction mixture need to be measured before any conclusions can be drawn on auto-ubiquitination from this experiment.

**Figure 5.**
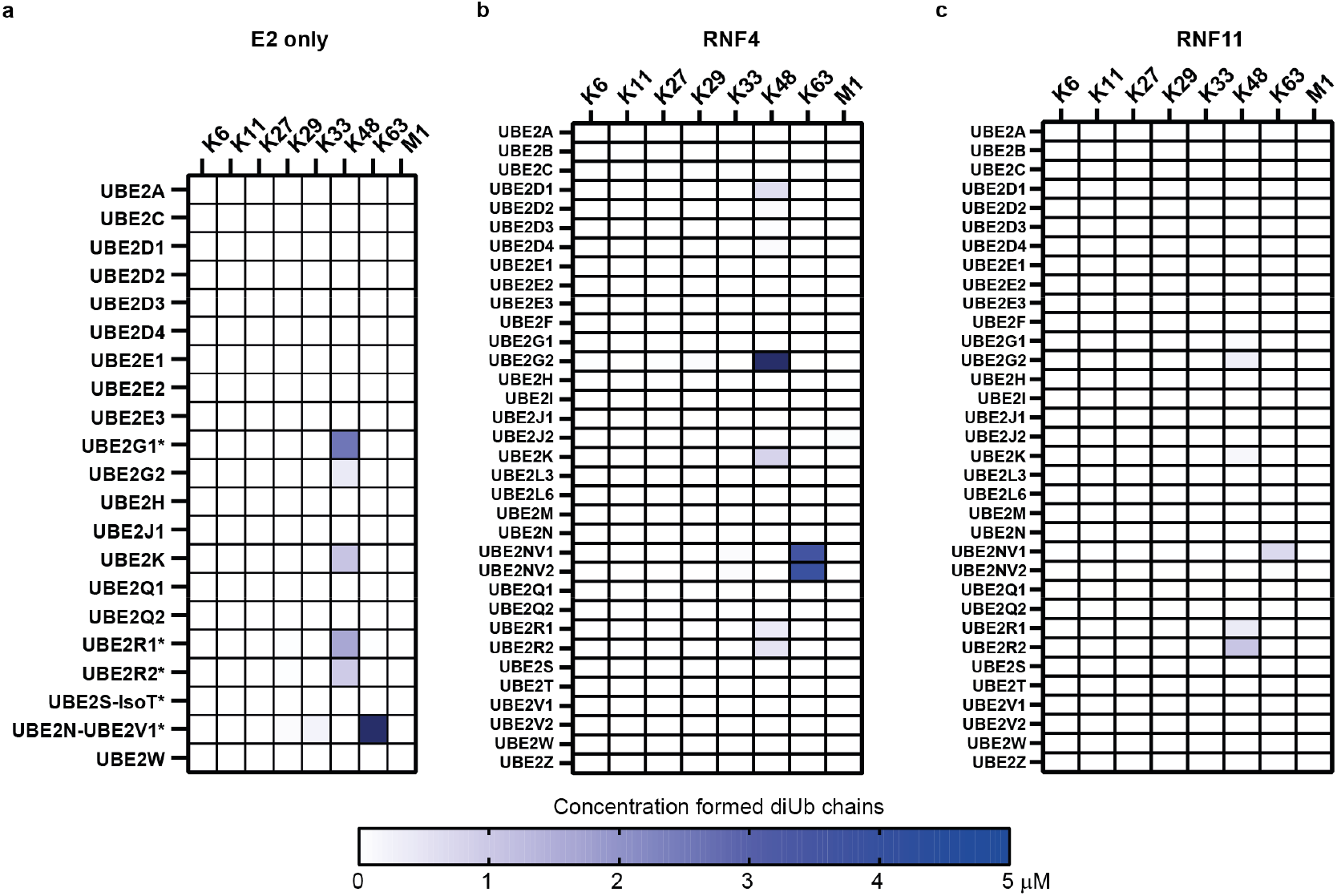
Assessment of *in vitro* diUb chain formation by different E2 enzymes and in combination with RNF4 or RNF11. **a**, E1 and E2 enzymes (100 nM and 1 μM, respectively) were incubated with the mixture of eight neutron-encoded monoUbs (5 μM each) and the donor Ub (40 μM) at 37°C for 24 hours. Reactions were stopped with 2% TFA and analyzed by mass spectrometry. DiUb chain formation was quantified. Asterisk (*) indicates that the E2 enzyme concentration is 2 μM instead of 1 μM. **b**,**c**, *In vitro* assay as described under **a**, but in the presence of 1 µM RNF4 (**b**) or RNF11 (**c**).

### Accelerating chain formation and determining active E2-E3 pairs

E3 ligase enzymes work in concert with an E1 and an E2 enzyme to deliver the activated ubiquitin onto its respective target. With at least 38 different E2 enzymes, 600-1000 different E3 enzymes, and many more potential substrates, there is an enormous amount of potential combinations. Therefore, convenient screening assays to identify active E2-E3 pairs and their products are highly desired. We used our set of neutron-encoded monoUbs to screen for the formation of diUb chains for 34 different recombinantly expressed E2 conjugation enzymes (**Supplementary Table 3**) in combination with two different E3 RING-ligases (RNF4 and RNF11) as a case study to prove that our developed multiplexed-MS assay can be used to screen active E2-E3 pairs and their products. (**Fig. 5b,c**). Overall, RNF4 enhanced the chain formation reaction of the E2 enzymes more than RNF11, and the amount of observed diUb chains is also higher for RNF4. With the help of RNF4, seven different E2 ligases showed diUb chain formation activity. UBED1, UBE2G2, UBE2K, UBE2R1 and UBE2R2 showed K48 chain formation. Especially the chain formation using UBE2G2 was significantly accelerated with the addition of RNF4 (compare **Fig. 5a** with **Fig. 5b**). Furthermore, UBE2N+UBE2V1 and UBE2N+UBE2V2 clearly built Lys63-linked chains. With the help of RNF11, Lys48-linked chain formation was observed for UBE2G1, UBE2G2, UBE2K, UBE2R1 and UBER2. UBE2N+UBE2V1 and UBE2N+UBE2V2 again built Lys63-linked chains, albeit at lower amount compared to RNF4.

## Discussion

We have developed a fast and sensitive multiplexed mass spectrometry assay to analyse the activity and selectivity of E2 enzymes and E2-E3 enzyme pairs *in vitro*, covering all diUb linkage types in a single measurement. It is important to gain more insight into these classes of enzymes since many of them are emerging as druggable targets.^23,41,42^ Moreover, these enzymes are becoming more attractive as key components of targeted protein degradation.^43^ Since much is unknown about their mode of action, activity and selectivity, a better understanding of the functions of these enzymes is important to further exploit them as drug targets and protein degraders.

Our assay can be used to screen for enzymatic activity and chain type selectivity *in vitro*. The key conceptual advantage of our MS assay lies in the ability to directly read out the type of chain that is built by an active E2/E3 pair in a single read-out. This was achieved by making use of synthetic monoUb substrates, which are modified to single-lysine mutants, with a ΔGlyGly C-terminal tail, and each mutant having its own fully ^13^C and ^15^N labelled amino acid fingerprint, resulting in eight monoUb isoforms with a distinguishable molecular weight. A non-isotope labelled, fully K→R Ub_1-76_ mutant was used as donor Ub to go through the enzymatic cascade. Depending on the lysine-side chain the donor Ub is conjugated to, a diUb chain with a distinguishable mass appears is produced. The effect of altering the substrate or enzyme concentration, or the influence of an allosteric binder or inhibitor, can easily be measured and quantified in a matter of minutes. The straightforward nature of the experimental set-up makes it possible to give an overview of the activity and selectivity of an E2-E3 pair in a single read-out without extensive sample preparation, and makes this assay attractive to use in other biochemical laboratories that are working with Ub conjugating and ligase enzymes.

Although some of the kinetic parameters are different and the actual native substrates are not used, the results to build selectively diUb chains with specific E2 enzymes and E2/E3 ligases were in accordance with literature. UBE2L3 together with NleL formed diUb chains from which two-thirds are linked via Lys6 and the other chains are connected via Lys48. These results are in accordance with literature, where predominantly Lys6-chain formation is observed, but also Lys48 is detected.^12,13^ Also, the observation that UBE2L3 in combination with UBE3C builds ∼17% Lys29-linked Ub chains and ∼80% Lys48-linked Ub is of a similar order as the results reported in literature.^22^ It is important to note here that both free Ub chains and Ub chains conjugated to the enzymes are taken into account in the analysis reported in literature, whereas in our case, we only consider diUb formation. The E3 ligase AREL1 in combination with UBE2L3 produced ∼88% Lys33-linked chains in our assay, where literature reported that the isolated triUb after chain assembly consisted of 75% Lys33-linked chains.^22^ However, AREL1 also should produce Lys11-linked diUb chains, which was not observed in our assay. The selective production of Lys48-linked chains by UBE2G1, UBE2R1 and UBE2R2 was in accordance with literature,^36,44^ and the same holds for the selective and efficient production of Lys63-linked chains by UBE2N-UBE2V1^45^ and the selective formation of Met1-linked chains by HOIP.^30^ However, UBE2S-IsoT was supposed to build Lys11-linked Ub chains, but no diUb chains were observed during our MS assay. The activity of UBE2S-IsoT was tested in an orthogonal assay with SDS-PAGE analysis as a read-out (**Supplementary Figure 7**). Ub chain formation and autoubiquitination of UBE2S-IsoT were observed on gel, meaning the enzyme was active. Ub chain formation between the Ub donor and the K11 acceptor-only was also examined. Here, autoubiquitination was observed, but no Ub chains were formed. Therefore, the substrate inhibition by other acceptors present in the MS assay could be excluded. However, for our assay, we used modified substrates. While it is known from literature that K→R mutations are accepted by UBE2S-IsoT,^29^ acetylation of different Lys positions, on the other hand, can inhibit Ub chain formation or hamper Ub elongation.^46^ Although this has not been specifically tested for N-terminal acetylation, it can potentially inhibit chain formation. Whether this modification can potentially influence the chain formation of other enzyme(s) (pairs) remains to be elucidated.

In addition to what is known in literature about these conjugation and ligation enzymes, our assay gave more insights into the relative speed of chain formation between the different enzymes and pairs. For example, we observed that the formation of Met1-linked diUb chains was much slower (24h) than the formation of Lys6- and Lys48-linked chains by UBE2L3+NleL (10 minutes). Furthermore, we found that chain formation by UBE2N+UBE2V1 and UBE2G2 was accelerated by the addition of RNF4, and we could analyse whether chain selectivity is enzyme concentration dependent, which was unlike DUBs^24,25^, not the case for the tested enzymes and within the applied concentration window.

We also used our assay to screen the ubiquitin chain-building capacity of E2 ligases. With this screen, we confirmed the chain conjugation capacity of six E2 conjugation enzymes. These conjugation capacities and the formed Ub chain types are known in literature.^36,40,45,47^ Unfortunately, no new insights in chain building were obtained from this screen. We also performed a proof-of-principle screen with a panel of 34 different E2 enzymes and two different RING-ligases to check whether they form an active conjugation pair and which chains they build. This screen resulted in seven active E2/RNF4 pairs and four less active E2/RNF11 pairs. Only Lys48- and Lys63-diUb formation was observed after 24h. The activity of UBE2G2 and UBE2D1 were enhanced by RNF4, and a larger amount of Lys48-linked diUb chains was produced. The activity of UBE2K and the UBE2N and UBE2V1 combination was not substantially enhanced by RNF4, and the amount of produced diUb chains for UBE2G1, UBE2R1 and UBE2R2 was lower for the combinations of these E2’s with RNF4 compared to the E2-only experiments. However, 2 μM of these E2 enzymes was present during the E2-only screen, while only 1 μM of these conjugation enzymes was used in the activity screen with RNF4, making direct comparison of the activity of RNF4 on the chain-building productivity difficult. So, RNF4 can accelerate chain formation in combination with some E2 ligases, however, not all E2 ligases benefit from it to the same extent. The amount of produced diUb chains of the screened E2 ligases in combination with RNF11 was lower in all cases in comparison with the E2-only screen. It is expected that although RNF11 may not enhance the production of diUb chains, at least a similar amount of diUb chains can be produced by the E2 enzymes present in the mixture. The lower amounts of diUb in the RNF11 screen might be explained by RNF11 binding diUb chains, which are not released in the reaction mixture or on the column of the LC-MS.

A limitation of our assay is the use of modified monoUb substrates that limit the scope to the formation of diUb chains. Some E2-(E3) ligases prefer to build longer chains or perform autoubiquitination, so if chain formation is not observed in our assay, it does not directly mean these enzymes do not build chains, like the UBE2D3 and TRIM25 combination. We can easily extend the scope of the assay into the field of building branched chains by making two lysine positions available on the acceptor Ub. In that case, the production of branched Ub chains could also be monitored with our assay. Furthermore, by changing the donor Ub for a donor Ub-like protein, the formation of heterologous chains can be measured.

The kinetics of our assay is different from normal kinetics due to the use of artificial substrates. A substrate with the incorrect Lys position available or the donor Ub can still bind the E2 or E3 enzyme, but the transfer of donor Ub onto this substrate is not possible, thereby effectively inhibiting the ligase activity. Dissociation of this substrate needs to happen before a new Ub can bind for possible transfer, so the stability of the E3∼Ub thioester must survive multiple binding- and dissociation cycles before a Ub chain could be built. Nevertheless, if the enzyme pair is able to form diUb chains, their activity and selectivity can be compared to other active enzyme pairs side-by-side.

The assay can be further optimized in the future. Currently, a lot of donor Ub needs to be present to be loaded onto the enzymes and to account for auto-ubiquitination. Since monoUb is more efficiently detected than diUb in the mass spectrometer, a very high starting signal of monoUb is observed. The concentration of monoUb which is loaded for the HPLC-MS analysis could also be further optimized. In combination with further gradient optimization, the broad peak of starting material can be transformed into a sharp peak over a small time window. Therewith, lower amounts of formed diUb are potentially easier to detect.

Another challenge of the designed assay is that during the time course, diUb is appearing, but sometimes after longer incubation times, the amount of measured diUb is decreasing. This observation can be caused by several reasons. If the formed Ub chain binds to the E2 or E3 enzyme and is not completely denatured, it cannot be observed in MS analysis. Another possibility is that during acidification of the sample, the enzymes will precipitate and the free Ub chains can (partly) do the same. Switching to Protein LoBind Tubes, a different acidification strategy and/or improved denaturing conditions could potentially solve this problem and therewith provide even clearer data on Ub chain assembly.

Besides decreasing diUb concentrations, sometimes diUb concentrations higher than 5 µM were measured, while with 5 µM of each acceptor monoUb present, only 5 µM of diUb can theoretically be formed. There are multiple potential explanations for this observation. The first is that the concentrations of the internal standard of the monoUb and diUb are different and therewith both signals are normalized to the same value, while this should be a different value. Another explanation is that the modified diUbs where 6 or 7 lysines are replaced by arginines ionize better and as such give higher signals for the same concentration of diUb in comparison with the internal standard. The third explanation could be that the broad signal of the monoUbs is overlapping in the chromatogram with the diUb signal(s), thereby influencing the detection efficiency of the diUbs present in the reaction mixture.

In conclusion, we report a novel mass spectrometry-based assay to screen for E2/E3 activity and selectivity for assembling diUb chains. This assay relies on synthetic modified neutron-encoded monoUbs, each having a distinguishable molecular weight, to allow for selective chain formation and direct detection and quantification of the assembled chain type. We showed that with our assay we could selectively build Ub chains with E2 enzymes and E2/E3 pairs known from literature. Furthermore, we showed that we could use our assay set-up to screen for E2 chain-building activity and for active E2/E3 pairs that have chain-building capacities. The assay has several advantages over existing methods: it requires low amounts of material, it is quantitative, the read-out can be done in a mid-throughput fashion, and a direct overview of activity and selectivity can be obtained from a single run. The easily adaptable assay set-up makes it possible to screen for the formation of branched Ub chains and heterologous chain formation in the future. Based on the proof-of-concept studies described here, we anticipate that our multiplex mass spectrometry-based assay will facilitate future discoveries on Ub chain assembly.

## Acknowledgements

This study is dedicated to the memory of Prof. Dr. Huib Ovaa, a great scientist, who passed away too soon on May 19^th^, 2020. We thank Alfred Vertegaal for the scientific discussions and Angela el Hebieshy for the expression and purification of the E2-E3 fusion protein gp78RING-Ube2g2 and purification of UBE1. We also thank Cami Talavera Ormeño and Paul Hekking for assistance with solid-phase peptide synthesis, and Angeliki Moutsiopoulou is thanked for her assistance with the expression and purification of recombinant ligases. pFlagCMV2-EFP was a gift from Dong-Er Zhang (Addgene plasmid # 12449), hUbc13 was a gift from Cynthia Wolberger (Addgene plasmid # 51131), and UEV1 was a gift from Cheryl Arrowsmith (Addgene plasmid # 25619). pMCSG17-NleL (Addgene plasmid #66716), pOPINS-AREL1 (Addgene plasmid #66710) and pOPINS-UBEC3 (Addgene plasmid #66711) were all kind gifts from David Komander.

This project is funded by the Institute for Chemical Immunology (grant no. ICI00026 to A.S.), by the TRIM-NET project, which received funding from the European Union’s Horizon 2020 research and innovation programme under the Marie Skłodowska-Curie grant agreement No 813599 to R.M., the Innovative Medicines Initiative 2 (IMI2) Joint Undertaking under grant agreement no. 875510 (EUbOPEN project) to B.R.v.D. and P.P.G, and by a VENI and VIDI grant (no. 722.014.002 and no. VI.Vidi.192.011) to G.J.v.d.H.v.N. from The Netherlands Organization for Scientific Research (NWO).

## Author Contributions

The concept of this study was designed by P.P.G. and B.D.M.v.T. Ub analogues were designed, synthesized and purified by B.D.M.v.T. Conjugation enzymes were expressed by B.D.M.v.T., J.L. and R.M. Biochemical assays were performed by B.D.M.v.T. Mass spectrometry support was done by B.R.v.D. Data analysis was performed by B.D.M.v.T with the assistance of G.S.M.L.K. Data analysis software was adjusted by B.C.J. Assistance with the choice of conjugation enzymes and the experimental design was given by J.L. and B.A.S. The manuscript was prepared by B.D.M.v.T. and P.P.G. with input from all authors. The project was supervised by P.P.G, G.J.v.d.H.v.N., and M.W.

## References

1. Wilkinson, K. D. & Audhya, T. K. Stimulation of ATP-dependent proteolysis requires ubiquitin with the COOH-terminal sequence Arg-Gly-Gly. J. Biol. Chem. 256, 9235–9241 (1981).

2. Komander, D. & Rape, M. The ubiquitin code. Annu. Rev. Biochem. 81, 203–229 (2012).

3. Tracz, M. & Bialek, W. Beyond K48 and K63: non-canonical protein ubiquitination. Cell. Mol. Biol. Lett. 26, 1–17 (2021).

4. Schulman, B. A. & Wade Harper, J. Ubiquitin-like protein activation by E1 enzymes: The apex for downstream signalling pathways. Nat. Rev. Mol. Cell Biol. 10, 319–331 (2009).

5. Ye, Y. & Rape, M. Building ubiquitin chains: E2 enzymes at work. Nat. Rev. Mol. Cell Biol. 10, 755– 764 (2009).

6. Buetow, L. & Huang, D. T. Structural insights into the catalysis and regulation of E3 ubiquitin ligases. Nat. Rev. Mol. Cell Biol. 17, 626–642 (2016).

7. Deshaies, R. J. & Joazeiro, C. A. P. RING domain E3 ubiquitin ligases. Annu. Rev. Biochem. 78, 399–434 (2009).

8. Li, W. et al. Genome-wide and functional annotation of human E3 ubiquitin ligases identifies MULAN, a mitochondrial E3 that regulates the organelle’s dynamics and signaling. PLoS One 3, 1–14 (2008).

9. Metzger, M. B., Hristova, V. A. & Weissman, A. M. HECT and RING finger families of E3 ubiquitin ligases at a glance. J. Cell Sci. 125, 531–537 (2012).

10. Walden, H. & Rittinger, K. RBR ligase-mediated ubiquitin transfer: A tale with many twists and turns. Nat. Struct. Mol. Biol. 25, 440–445 (2018).

11. Popovic, D., Vucic, D. & Dikic, I. Ubiquitination in disease pathogenesis and treatment. Nat. Med. 20, 1242–1253 (2014).

12. Hospenthal, M. K., Freund, S. M. V. & Komander, D. Assembly, analysis and architecture of atypical ubiquitin chains. Nat. Struct. Mol. Biol. 20, 555–565 (2013).

13. Lin, D. Y., Diao, J., Zhou, D. & Chen, J. Biochemical and Structural Studies of a HECT-like Ubiquitin Ligase from Escherichia coli O157:H7. J. Biol. Chem. 286, 441–449 (2011).

14. Hospenthal, M. K., Mevissen, T. E. T. & Komander, D. Deubiquitinase-based analysis of ubiquitin chain architecture using Ubiquitin Chain Restriction (UbiCRest). Nat. Protoc. 10, 349–361 (2015).

15. Kirkpatrick, D. S. et al. Quantitative analysis of in vitro ubiquitinated cyclin B1 reveals complex chain topology. Nat. Cell Biol. 8, 700–710 (2006).

16. Meierhofer, D., Wang, X., Huang, L. & Kaiser, P. Quantitative analysis of global ubiquitination in HeLa cells by mass spectrometry. J. Proteome Res. 7, 4566–4576 (2008).

17. Phu, L. et al. Improved quantitative mass spectrometry methods for characterizing complex ubiquitin signals. Mol. Cell. Proteomics 10, 1–19 (2011).

18. Peng, J. et al. A proteomics approach to understanding protein ubiquitination. Nat. Biotechnol. 21, 921–926 (2003).

19. Kim, W. et al. Systematic and quantitative assessment of the ubiquitin-modified proteome. Mol. Cell 44, 325–340 (2011).

20. Hjerpe, R. et al. Efficient protection and isolation of ubiquitylated proteins using tandem ubiquitin-binding entities. EMBO Reports 10, 1250–1258 (2009).

21. Swatek, K. N. et al. Insights into ubiquitin chain architecture using Ub-clipping. Nature 572, 533– 537 (2019).

22. Michel, M. A. et al. Assembly and Specific Recognition of K29- and K33-Linked Polyubiquitin. Mol. Cell 58, 95–109 (2015).

23. De Cesare, V. et al. The MALDI-TOF E2/E3 Ligase Assay as Universal Tool for Drug Discovery in the Ubiquitin Pathway. Cell Chem. Biol. 25, 1117–1127 (2018).

24. Mevissen, T. E. T. et al. OTU Deubiquitinases Reveal Mechanisms of Linkage Specificity and Enable Ubiquitin Chain Restriction Analysis. Cell 154, 169–184 (2013).

25. Ritorto, M. S. et al. Screening of DUB activity and specificity by MALDI-TOF mass spectrometry. Nat. Commun. 5, 4763 (2014).

26. Blythe, E. E., Olson, K. C., Chau, V. & Deshaies, R. J. Ubiquitin-A nd ATP-dependent unfoldase activity of P97/VCP•NPLOC4•UFD1L is enhanced by a mutation that causes multisystem proteinopathy. Proc. Natl. Acad. Sci. U. S. A. 114, E4380–E4388 (2017).

27. Pickart, C. M. & Raasi, S. Controlled synthesis of polyubiquitin chains. Methods Enzymol. 399, 21– 36 (2005).

28. Michel, M. A., Komander, D. & Elliott, P. R. Enzymatic assembly of ubiquitin chains. Methods Mol. Biol. 1844, 73–84 (2018).

29. Bremm, A., Freund, S. M. V. & Komander, D. Lys11-linked ubiquitin chains adopt compact conformations and are preferentially hydrolyzed by the deubiquitinase Cezanne. Nat. Struct. Mol. Biol. 17, 939–947 (2010).

30. Dong, K. C. et al. Preparation of distinct ubiquitin chain reagents of high purity and yield. Structure 19, 1053–1063 (2011).

31. Jansen, B. C. et al. LaCyTools: A Targeted Liquid Chromatography-Mass Spectrometry Data Processing Package for Relative Quantitation of Glycopeptides. J. Proteome Res. 15, 2198–2210 (2016).

32. Falck, D., Jansen, B. C., de Haan, N. & Wuhrer, M. High-throughput analysis of IgG Fc glycopeptides by LC-MS. Methods Mol. Biol. 1503, 31–47 (2017).

33. Choudhury, N. R. et al. RNA-binding activity of TRIM25 is mediated by its PRY/SPRY domain and is required for ubiquitination. BMC Biol. 15, 1–20 (2017).

34. Koliopoulos, M. G., Esposito, D., Christodoulou, E., Taylor, I. A. & Rittinger, K. Functional role of TRIM E3 ligase oligomerization and regulation of catalytic activity. EMBO J. 35, 1204–1218 (2016).

35. Marblestone, J. G. et al. Comprehensive Ubiquitin E2 Profiling of Ten Ubiquitin E3 Ligases. Cell Biochem. Biophys. 67, 161–167 (2013).

36. Sheng, Y. et al. A human ubiquitin conjugating enzyme (E2)-HECT E3 ligase structure-function screen. Mol. Cell. Proteomics 11, 329–341 (2012).

37. Stewart, M. D., Ritterhoff, T., Klevit, R. E. & Brzovic, P. S. E2 enzymes: More than just middle men. Cell Res. 26, 423–440 (2016).

38. Pickart, C. M. & Rose, I. A. Functional heterogeneity of ubiquitin carrier proteins. J. Biol. Chem. 260, 1573–1581 (1985).

39. Haldeman, M. T., Xia, G., Kasperek, E. M. & Pickart, C. M. Structure and function of ubiquitin conjugating enzyme E2-25K: The tail is a core-dependent activity element. Biochemistry 36, 10526–10537 (1997).

40. David, Y., Ziv, T., Admon, A. & Navon, A. The E2 ubiquitin-conjugating enzymes direct polyubiquitination to preferred lysines. J. Biol. Chem. 285, 8595–8604 (2010).

41. Bielskiene, K., Bagdoniene, L., Mozuraitiene, J., Kazbariene, B. & Janulionis, E. E3 ubiquitin ligases as drug targets and prognostic biomarkers in melanoma. Med. 51, 1–9 (2015).

42. Rossi, M. et al. High-throughput screening for inhibitors of the HECT ubiquitin E3 ligase ITCH identifies antidepressant drugs as regulators of autophagy. Cell Death Dis. 5, 1–12 (2014).

43. Gross, P. et al. Accelerating PROTAC drug discovery: Establishing a relationship between ubiquitination and target protein degradation. Biochem. Biophys. Res. Commun. 628, (2022).

44. David, Y., Ziv, T., Admon, A. & Navon, A. The E2 ubiquitin-conjugating enzymes direct polyubiquitination to preferred lysines. J. Biol. Chem. 285, 8595–8604 (2010).

45. Petroski, M. D. et al. Substrate modification with lysine 63-linked ubiquitin chains through the UBC13-UEV1A ubiquitin-conjugating enzyme. J. Biol. Chem. 282, 29936–29945 (2007).

46. Ohtake, F. et al. Ubiquitin acetylation inhibits polyubiquitin chain elongation. EMBO Rep. 16, 192– 201 (2015).

47. Chen, Z. & Pickart, C. M. A 25-kilodalton ubiquitin carrier protein (E2) catalyzes multi-ubiquitin chain synthesis via lysine 48 of ubiquitin. J. Biol. Chem. 265, 21835–21842 (1990).

48. Van Tol, B.D.M. et al. Neutron-encoded diubiquitins to profile linkage selectivity of deubiquitinating enzymes. Nat. Commun. 14, 1661 (2023).

