## Supplementary Information for "Employing neutron-encoded monoUbs to study E2/E3 ligase activity and selectivity for assembling Ub chains"

#### Materials and Methods

#### Table of contents

**Supplementary Figures, Schemes and Tables**
Supplementary Figure 1. Full gel image of all eight neutron-encoded acceptor monoUbs and donor monoUb
Supplementary Figure 2. Full MS spectrum of the mix of all eight neutron-encoded acceptor monoUbs and donor monoUb
Supplementary Figure 3. Full gel images of proof of principle assay of Figure 3
Supplementary Figure 4. Determination of the assembly of all different diUb linkages built by specific E2 conjugating enzymes and E2-E3 pairs.
Supplementary Figure 5. Determination of the assembly of diUb linkages built by specific E2 conjugating enzymes combined with different RING ligases.
Supplementary Figure 6. IsoT activity analysis.
Supplementary Scheme 1. Synthesis of all nine neutron-encoded monoUbs
Supplementary Table 1. Amino acid sequence of the designed monoubiquitins
Supplementary Table 2. Purified recombinant ligase enzymes used in this work.
Supplementary Table 3. E2 enzymes used for RING ligase screening
Supplementary Table 4. Alignment file/table
Supplementary Table 5. LaCyTools settings
Supplementary Table 6. Analytes and calibrants – LaCyTools

**Materials and Methods; Protein Synthesis**
General
Solid Phase Peptide Synthesis
General procedures, analysis and purification
Synthesis of neutron-encoded monoubiquitins
Synthesis of internal standards

**Materials and Methods; Biochemistry**
Recombinant protein expression and purification
General method SDS-PAGE analysis
SDS-PAGE analysis:
* Characterization of all eight neutron-encoded monoUbs, donor monoUb, K48 click diUb and $Ub_{1-74}$
* E1-E2-E3 chain building capacity of modified substrates
* IsoT activity analysis
*In vitro* conjugation assays with mass spectrometry read-out

**References**

#### Supplementary Figures, Schemes and Tables

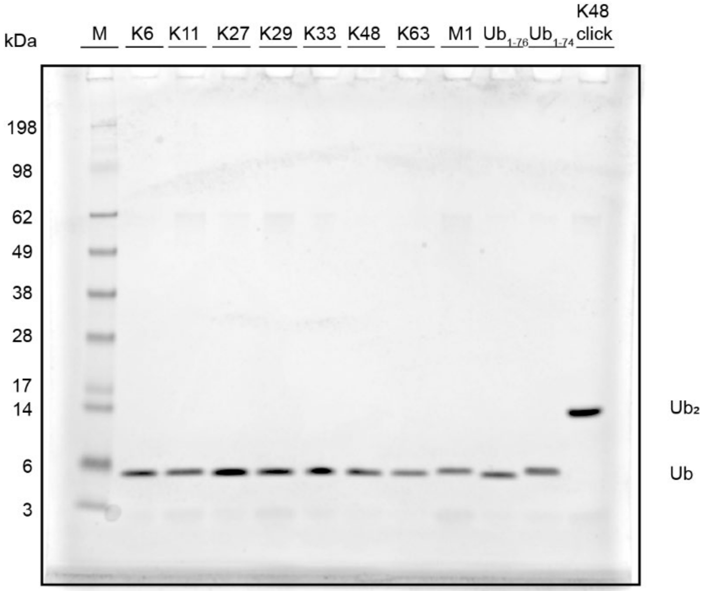


**Supplementary Figure 1| Full gel image of all eight neutron-encoded acceptor monoUbs (4a-g and 5), donor monoUb (6), non-hydrolysable clicked Lys48 diUb and $Ub_{1\text{-}74}$ on 12% Bis-Tris gel related to Figure 2c in the main paper.**

Protein marker = SeeBlue™ Plus2 Pre-stained Protein Standard. Loading: ~0.62 µg/lane.
*Method*; Stock solutions of all Ubs were diluted in a buffer containing 20 mM TRIS and 200 mM NaCl to a concentration of ~3.5 µM. 5 µL of sample buffer was added to 10 µL of diluted stock solution. The samples were boiled for 5 min at 95°C and 7.5 µL was loaded on a precast 4-12%Bis-Tris gel (Invitrogen). The samples were resolved by gel electrophoresis with MES running buffer and the gel was stained with InstantBlue™ staining.

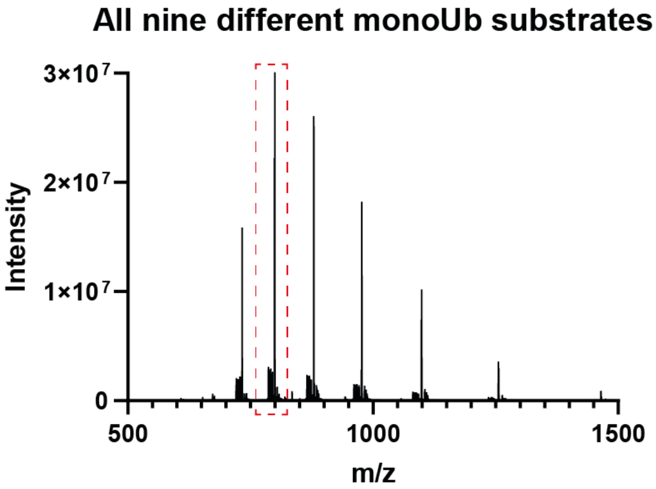


**Supplementary Figure 2| Sum spectrum of the main peak of the chromatogram of the mix of all eight neutron-encoded monoUbs (5 µM each) and donor monoUb (40 µM). Full mass range of sum spectrum of Figure 2d in the main paper.**

*Method;* All eight neutron-encoded monoUbs were mixed in equimolar amount (Final concentration = 5 µM for each monoUb) and one equivalent of donor Ub was added for each acceptor monoUb (Final concentration = 40 µM). This mixture was 24 times diluted with 0.1% FA in MQ and 4 µL of this solution was injected for a HPLC-MS run (LC-MS – System 2 – Gradient 1). Spectra 103:153 were combined for the sum spectrum. Zoom in of Figure 2d is from the area within the red dotted box.

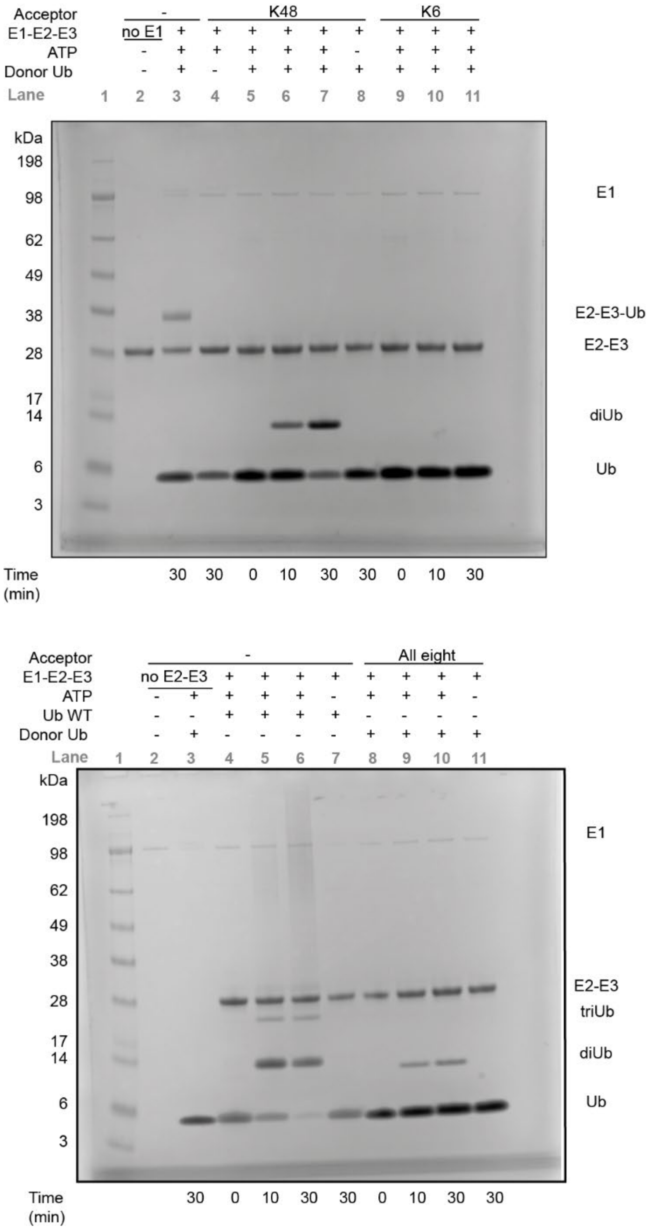


**Supplementary Figure 3| Full gel images of proof of principle assay on 4-12% Bis-Tris gel related to Figure 3 in the main paper.**
Protein marker = SeeBlue™ Plus2 Pre-stained Protein Standard. Loading: 5 µL reaction mixture.
*Method*; Quenched samples of the reaction mixtures were boiled for 5 min. at 95°C and 7.5 µL was loaded on gel. The samples were resolved by gel electrophoresis on a 4-12% Bis-Tris gel (Invitrogen) with MES running buffer and the gels were stained with InstantBlue™ staining.

[Figure 4 is included at the end of this document]

**Supplementary Figure 4| Determination of the assembly of all different diUb linkages built by specific E2 conjugating enzymes and E2-E3 pairs**. The quantified assay results are plotted. The first graph shows the amount of diUb (μM) formed in the reaction mixture normalized to the internal standard non-hydrolysable Lys48-linked diUb and calculated using the concentration of the internal standard as reference. The second graph shows the amount of monoUb (μM) present in the reaction mixture, normalized to the internal standard $Ub_{1-74}$ and calculated using the concentration of the internal standard as reference.

[Figure 5 is included at the end of this document]

**Supplementary Figure 5| Determination of the assembly of diUb linkages built by specific E2 conjugating enzymes combined with different RING ligases**. The quantified assay results are plotted. The first graph shows the amount of diUb (μM) formed in the reaction mixture normalized to the internal standard non-hydrolysable Lys48-linked diUb and calculated using the concentration of the internal standard as reference. The second graph shows the amount of monoUb (μM) present in the reaction mixture, normalized to the internal standard $Ub_{1-74}$ and calculated using the concentration of the internal standard as reference.

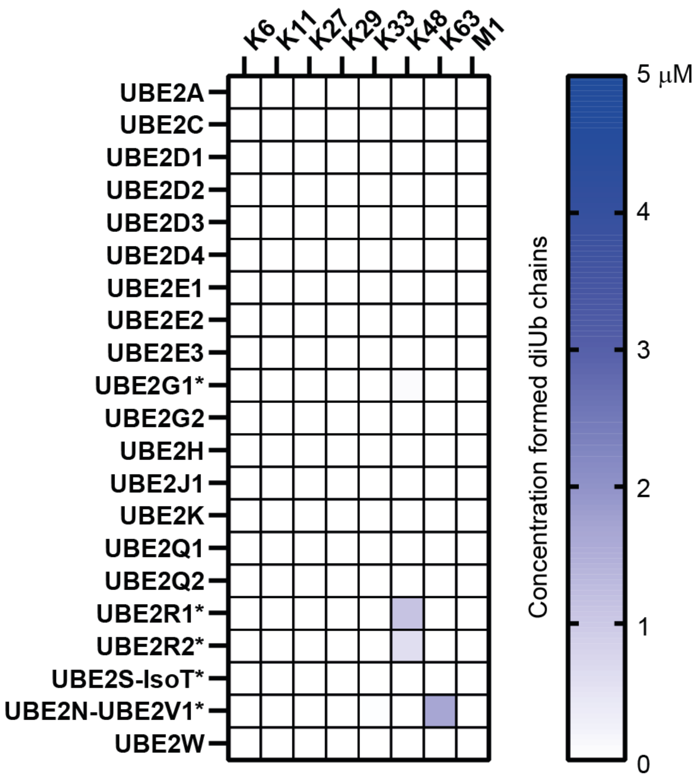


**Supplementary Figure 6| *In vitro* chain formation by E2 enzymes.** E1 and E2 enzymes (100 nM and 1 μM, respectively) were incubated with the mixture of eight neutron-encoded monoUbs (5 μM each) and the donor Ub (40 μM) at 37°C for 1 hour. Reactions were stopped wit 2% TFA and analyzed by mass spectrometry. DiUb chain formation was quantified.

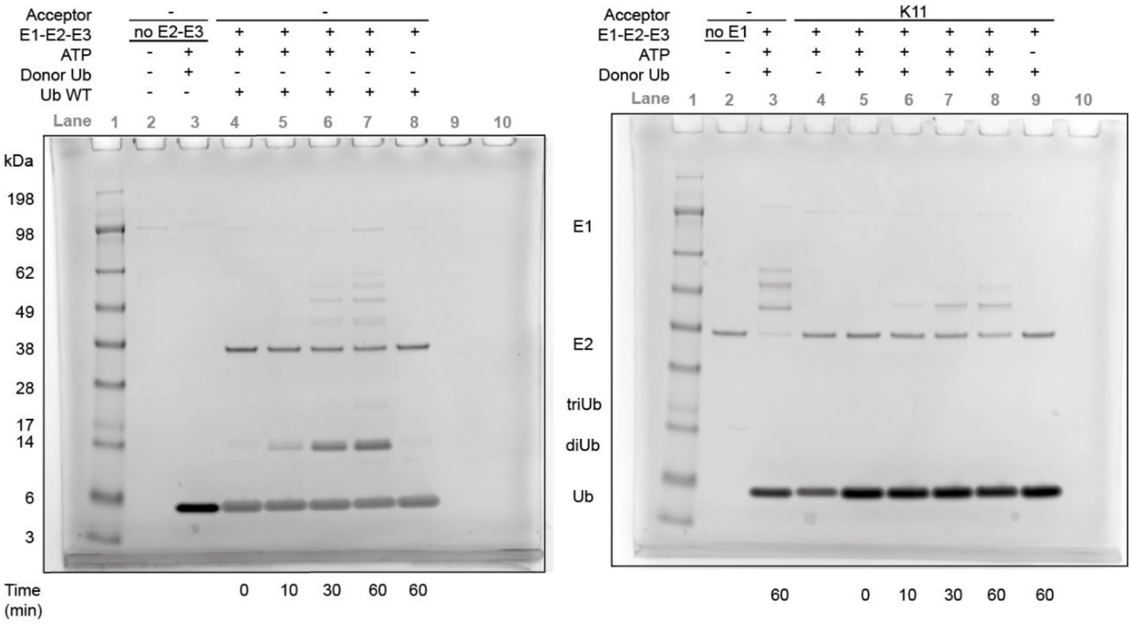


**Supplementary Figure 7| IsoT activity analysis.**

Protein marker = SeeBlue™ Plus2 Pre-stained Protein Standard. Loading: 5 μL reaction mixture.

*Method*; Quenched samples of the reaction mixtures were boiled for 5 min. at 95°C and 7.5 μL was loaded on gel. The samples were resolved by gel electrophoresis on a 4-12% Bis-Tris gel (Invitrogen) with MES running buffer and the gels were stained with InstantBlue™ staining.

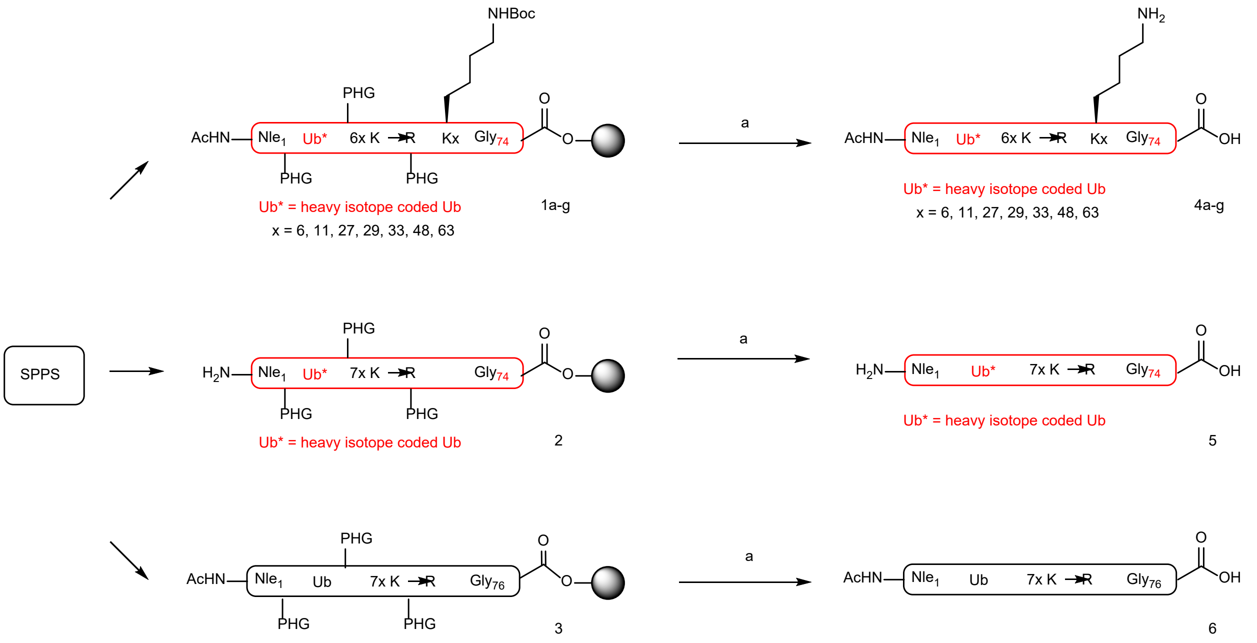


**Supplementary Scheme 1| Synthesis of all nine neutron-encoded monoUbs.** Reagents and conditions: (a) TFA/$H_2O$/PhOH/*i*$Pr_3SiH$ (90.5/5/2.5/2; v/v/v/v). Monoubiquitins **1a-g**, **2** and **3** on resin were synthesized using solid phase peptide synthesis (SPPS). Monoubiquitins **1a-g, 2** and **3** were liberated from the resin and deprotected using 90% TFA, yielding neutron-encoded $Ub_{1-74}$ containing 6x K→R mutation and an acetylated N-terminus **4a-g**, neutron-encoded $Ub_{1-74}$ containing 7x K→R mutation and a free N-terminus **5** and $Ub_{1-76}$ containing 7x K→R mutation and an acetylated N-terminus **6**.

**Supplementary Table 1| Amino acid sequence of the designed monoubiquitins.** Sites of neutron-encoded amino acids, acetyl groups, pseudoproline building blocks and dipeptides incorporation and Lys to Arg mutations are indicated and relevant structures are shown below.

| Ubiquitin Linkage | Protein sequence for synthesizer | Amount of neutron-encoded amino acids | Mass difference in comparison with unlabeled ubiquitin | Mass difference in comparison with previous ubiquitin | Molecular Weight |
|---|---|---|---|---|---|
| Donor Ub | AcNleQIFVRTLTGRTITLEVEPSDTIENVRARIQDREGIPPDQQRLIFAGRQLEDGRTLSDYNIQRESTLHLVLRLRGG | | | | 8784,97 Da |
| K6 | AcNleQIFVKTLTGRTITLEVEPSDTIENVRARIQDREGIPPDQQRLIFAGRQLEDGRTLSDYNIQRESTLHLVLRLR | | 0 Da | - | 8642,85 Da |
| K11 | AcNleQIFVRTLTGKTITLEVEPSDTIENVRARIQDREGIPPDQQRLIFAGRQLEDGRTLSDYNIQRESTLHLVLRLR | 1xV + 1xL | +13 Da | 13 Da | 8655,75 Da |
| K27 | AcNleQIFVRTLTGRTITLEVEPSDTIENVKARIQDREGIPPDQQRLIFAGRQLEDGRTLSDYNIQRESTLHLVLRLR | 3xV + 1xI | + 25 Da | 12 Da | 8667,66 Da |
| K29 | AcNleQIFVRTLTGRTITLEVEPSDTIENVRAKIQDREGIPPDQQRLIFAGRQLEDGRTLSDYNIQRESTLHLVLRLR | 3xV + 2xL + 1xI | + 39 Da | 14 Da | 8681,55 Da |
| K33 | AcNleQIFVRTLTGRTITLEVEPSDTIENVRARIQDKEGIPPDQQRLIFAGRQLEDGRTLSDYNIQRESTLHLVLRLR | 3xV + 2xL + 3xI | +53 Da | 14 Da | 8695,44 Da |
| K48 | AcNleQIFVRTLTGRTITLEVEPSDTIENVRARIQDREGIPPDQQRLIFAGKQLEDGRTLSDYNIQRESTLHLVLRLR | 3xV + 4xL + 3xI | + 67 Da | 14 Da | 8709,33 Da |
| K63 | AcNleQIFVRTLTGRTITLEVEPSDTIENVRARIQDREGIPPDQQRLIFAGRQLEDGRTLSDYNIQKESTLHLVLRLR | 3xV + 4xL + 5xI | + 81 Da | 14 Da | 8723,22 Da |
| M1 | NleQIFVRTLTGRTITLEVEPSDTIENVRARIQDREGIPPDQQRLIFAGRQLEDGRTLSDYNIQRESTLHLVLRLR | 4xV + 6xL + 6xI | + 94 Da | 13 Da | 8736,04 Da |

Ac = Acetyl    Nle = NorLeucine    XX = dipeptide building block

X = neutron-encoded amino acid    R = Lys→Arg mutation

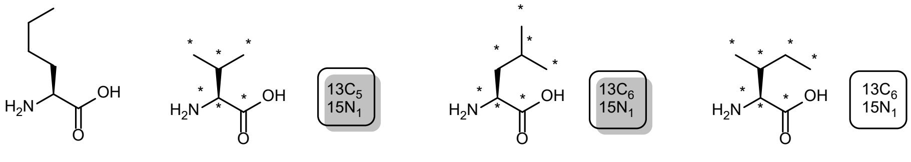


NorLeucine

Isotope labelled Valine
$13C_5H_{1115}NO_2$
mw: 123,08 g/mol

Isotope labelled Leucine
$13C_6H_{1315}NO_2$
mw: 138,09 g/mol

Isotope labelled Isoleucine
$13C_6H_{1315}NO_2$
mw: 138,09 g/mol

**Supplementary Table 2| Purified recombinant ligase enzymes used in this work.**

| | Ligase family | Human gene name | Domain/ Length / Fragment | Tag | UniProt accession number | Species / Origin / Organism | Expression system / Host / Source | Stock concentration | Source or reference |
|---|---|---|---|---|---|---|---|---|---|
| 1 | E1 ligase | Ube1 | unknown | N-terminal 6-His tagged | P22314 | Human | E. coli BL21(DE3) | 33.9 μM | *In house.* [Mulder, 2016][1] |
| 2 | E2 Family 4 | Ube2D1 | Full length | unknown | P51668 | Human | unknown | 100 μM | Gift from Brenda Schulman |
| 3 | E2 Family 4 | Ube2D2 | Full length | GST- tagged, clvd | P62837 | Human | E.coli BL21-Gold(DE3) | 100 μM | Gift from Brenda Schulman, [Liwocha, 2020][2] |
| 4 | E2 Family 4 | Ube2D3 | Full length | GST- tagged, clvd | P61077 | Human | E.coli BL21-Gold(DE3) | 100 μM | Gift from Brenda Schulman, [Liwocha, 2020][2] |
| 5 | E2 Family 3 | Ube2G1 | GS-FL(1-70) | N-terminal His- tagged, clvd | P62253 | Human | E.coli BL21-Gold(DE3) | 200 μM | Gift from Brenda Schulman, [Liwocha, 2020][2] |
| 6 | E2 Family 15 | Ube2L3 | Full length | unknown | P68036 | Human | unknown | 820 μM | Gift from Brenda Schulman |
| 7 | E2 Family 9 | Ube2N | Full length | His-tagged, clvd | P61088 | Human | E. coli BL21(DE3) | 49.2 μM | *In house,* This work |
| 8 | E2 Family 3 | Ube2R1 | GS-FL(1-236) | unknown | P49427 | Human | unknown | 80 μM | Gift from Brenda Schulman |
| 9 | E2 Family 3 | Ube2R2 | GS-FL(1-238) | GST- tagged, clvd | Q172K3 | Human | unknown | 90 μM | Gift from Brenda Schulman, [Liwocha, 2020][2] |
| 10 | E2 Family 11 | Ube2S-IsoT | FL(1-196) fused with USP5/IsoT (res. 173-289) | GST- tagged, clvd | N/A | Human | E.coli BL21-Gold(DE3) | 400 μM | Gift from Brenda Schulman, [Liwocha, 2020][2] |
| 11 | E2 Family 10 | Ube2V1 | 8-147 | His-tagged, clvd | Q13404 | Human | E. coli BL21(DE3) | 97.7 μM | *In house,* This work |
| 12 | E2-E3 ligase fusion protein | Ube2G2-gp78 | Gp78/AMFR (322-398) fused to N-terminus of FL(1-165) UBE2G2 with GTGSH link | His-tagged, clvd | N/A | unknown | E. coli BL21(DE3) | 268.7 μM | *In house, [El Hiebeshy, 2023]* |
| 13 | Mimics eukaryotic HECT E3 ligase | NleL | 170-782 | GST- tagged, clvd | unkown | E.coli O157:H7 | E. coli BL21(DE3) | 279 μM | *In house,* This work |
| 14 | HECT E3 ligase | AREL1 | 436-823 | His-tagged, clvd | O15033 | Human | E. coli BL21(DE3) | 33 μM | *In house,* This work |
| 15 | HECT E3 ligase | UBE3C | 693-1083 | His-tagged, clvd | Q15386 | Human | E. coli BL21(DE3) | 327 μM | *In house,* This work |
| 16 | RING E3 ligase | RNF4 | RING-RING domain (134-194)/(130-190) | His-tagged, clvd | P78317 | unknown | E.coli Rosetta (DE3) | 420 μM | Gift from Brenda Schulman |
| 17 | RING E3 ligase | TRIM25 | Full length | GST- tagged, clvd | Q14258 | Human | E. coli BL21(DE3) | 70.9 μM | *In house,* This work |
| 18 | RBR E3 ligase | HOIP | 696-1072 | unknown | Q96EP0 | unknown | unknown | 71 μM | Gift from Brenda Schulman |

**Supplementary Table 3| E2 enzymes used for RING ligase screening**

| E2 conjugating Enzyme | Alternate Name | Tag |
|---|---|---|
| **Ube2A** | HR6A | No tag |
| **Ube2B** | HR6B | No tag |
| **Ube2C** | UBcH10 | T7 |
| **Ube2D1** | UbcH5A | T7 |
| **Ube2D2** | UbcH5B | T7 |
| **Ube2D3** | UbcH5C | No tag |
| **Ube2D4** | UbcH5D | T7 |
| **Ube2E1** | UbcH6 | No tag |
| **Ube2E2** | UbcH8 | T7 |
| **Ube2E3** | UbcH9 | No tag |
| **Ube2F** | NCE2 | T7 |
| **Ube2G1** | Ubc7 | T7 |
| **Ube2G2** | Ubc7 | No tag |
| **Ube2H** | UbcH2 | No tag |
| **Ube2I** | Ubc9 | No tag |
| **Ube2J1** | NCUBE1 | His-T7 |
| **Ube2J2** | NCUBE2 | T7 |
| **Ube2K** | Ubc1 | No tag |
| **Ube2L3** | UbcH7 | No tag |
| **Ube2L6** | UbcH8 | No tag |
| **Ube2M** | Ubc12 | No tag |
| **Ube2N** | Ubc13 | No tag |
| **Ube2N/Ube2V1** | Ubc13/Uev1A | No tag/T7 |
| **Ube2N/Ube2V2** | Ubc13/Mms2 | No tag/No tag |
| **Ube2Q** | NiCE-5 | His-T7 |
| **Ube2Q2** | - | No tag |
| **Ube2R1** | CDC34 | T7 |
| **Ube2R2** | CDC34B | T7 |
| **Ube2S** | E2-EPF | T7 |
| **Ube2T** | HSPC10 | No tag |
| **Ube2V1** | Uev1A | T7 |
| **Ube2V2** | Mms2 | No tag |
| **Ube2W** | Ubc16 | His-T7 |
| **Ube2Z** | USE1 | His-T7 |

*****Supplier Ubiquigent E2$^{scan™}$ Kit . Cat#67-0005-001

**Supplementary Table 4| Alignment file/table**

| *m/z* | **Seconds** |
|---|---|
| 599.3889 | 200 |
| 603.3051 | 199 |
| 616.0513 | 205 |
| 767.6136 | 209 |
| 795.0938 | 209 |
| 947.0273 | 212 |
| 799.5303 | 215 |
| 788.8981 | 216 |

**Supplementary Table 5| LaCyTools settings**

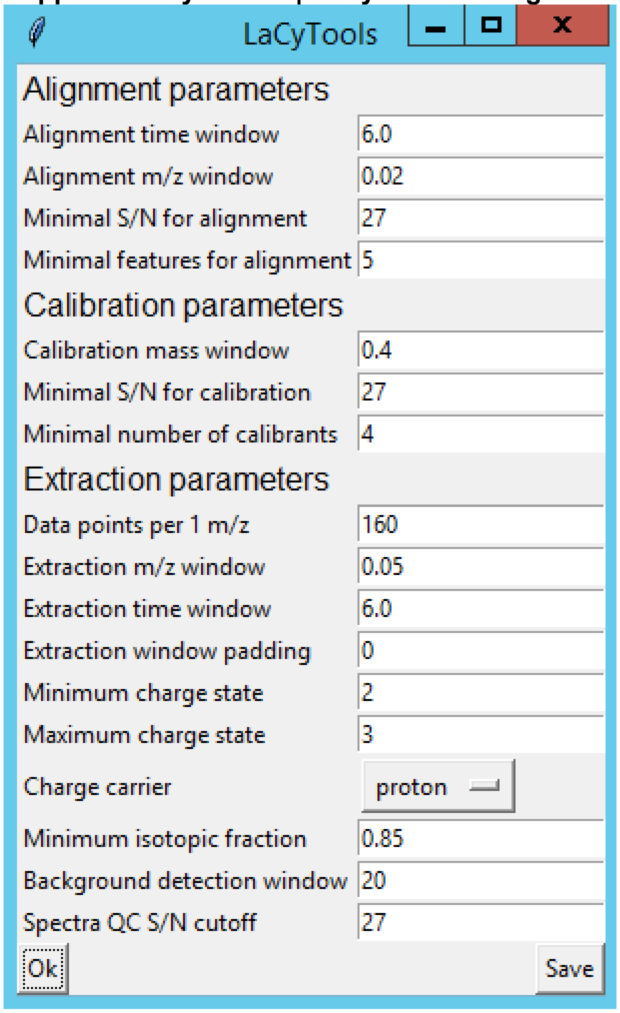

LaCyTools

##### Alignment parameters

| | |
|---|---|
| Alignment time window | 6.0 |
| Alignment m/z window | 0.02 |
| Minimal S/N for alignment | 27 |
| Minimal features for alignment | 5 |

##### Calibration parameters

| | |
|---|---|
| Calibration mass window | 0.4 |
| Minimal S/N for calibration | 27 |
| Minimal number of calibrants | 4 |

##### Extraction parameters

| | |
|---|---|
| Data points per 1 m/z | 160 |
| Extraction m/z window | 0.05 |
| Extraction time window | 6.0 |
| Extraction window padding | 0 |
| Minimum charge state | 2 |
| Maximum charge state | 3 |
| Charge carrier | proton |
| Minimum isotopic fraction | 0.85 |
| Background detection window | 20 |
| Spectra QC S/N cutoff | 27 |

Ok Save

**Supplementary Table 6| Analytes and calibrants – LaCyTools**

| Peak | RT | Mass Window | Time Window | Minimal charge state | Maximum charge state | Calibrant |
|---|---|---|---|---|---|---|
| _C756_H1255_N211_O233 | 213 | | | 10 | 25 | X |
| _C758_H1258_N234_O235 | 226 | | | 10 | 25 | X |
| _C747_hC11_H1258_N232_hN2_O235 | 226 | | | 10 | 25 | X |
| _C737_hC21_H1258_N230_hN4_O235 | 226 | | | 10 | 25 | X |
| _C725_hC33_H1258_N228_hN6_O235 | 226 | | | 10 | 25 | X |
| _C713_hC45_H1258_N226_hN8_O235 | 226 | | | 10 | 25 | X |
| _C701_hC57_H1258_N224_hN10_O235 | 226 | | | 10 | 25 | X |
| _C689_hC69_H1258_N222_hN12_O235 | 226 | | | 10 | 25 | X |
| _C664_hC92_H1256_N220_hN16_O234 | 226 | | | 10 | 25 | X |
| _C375_H625_N103_O116 | 209 | | | 5 | 13 | X |
| _C381_H633_N119_O119 | 215 | | | 5 | 13 | X |
| _C377_H627_N115_O117 | 215 | | | 5 | 13 | X |
| _C366_hC11_H627_N113_hN2_O117 | 215 | | | 5 | 13 | X |
| _C356_hC21_H627_N111_hN4_O117 | 215 | | | 5 | 13 | X |
| _C344_hC33_H627_N109_hN6_O117 | 215 | | | 5 | 13 | X |
| _C332_hC45_H627_N107_hN8_O117 | 215 | | | 5 | 13 | X |
| _C320_hC57_H627_N105_hN10_O117 | 215 | | | 5 | 13 | X |
| _C308_hC69_H627_N103_hN12_O117 | 215 | | | 5 | 13 | X |
| _C283_hC92_H625_N101_hN16_O116 | 209 | | | 5 | 13 | X |

_C = carbon atom = 12.00000
_hC = heavy carbon atom = 13.0033550
_H = hydrogen atom = 1.007825
_N = nitrogen atom = 14.003074
_hN = heavy nitrogen atom = 15.000109
_O = oxygen atom = 15.994915

#### Materials and Methods; Protein Synthesis

*General.* Solvents and reagents for peptide synthesis were purchased from various suppliers (listed in Table 7) and were used as received. Linear solid phase peptide synthesis of Ub was performed on resin according to an established method described by our group.[3] LC-MS data processing was performed using Waters MassLynx Mass Spectrometry Software 4.2. Deconvoluted mass was obtained from the electrospray ionization mass spectrum envelope (average isotopes) with Maxent1 function. The calculated mass of Ub (derivatives) was obtained with ChemDraw Professional 20.0 (PerkinElmer Informatics, Inc.) by calculating the molecular weight of the complete structure.

**Supplementary Table 7| Building blocks, reagents and solvents for ubiquitin synthesis.***

| Compound | Abbreviation | CAS# | Source or reference |
|---|---|---|---|
| ***Building blocks*** | | | |
| **Fmoc Gly TentaGel® R HMPA resin** | Gly-HMPA | - | Rapp Polymere GmbH, # 1513 |
| **Fmoc Arg(pbf) TentaGel® R HMPA resin** | Arg-HMPA | - | Rapp Polymere GmbH, # 1502 |
| **Fmoc-L-Valine-OH 13C5,15N1** | Fmoc-L-Val $^{13}C_5$ $^{15}N_1$ | - | In-house synthesis, **Chapter 2** |
| **Fmoc-L-Isoleucine-OH 13C6,15N1** | Fmoc-L-Ile $^{13}C_6$ $^{15}N_1$ | - | In-house synthesis, **Chapter 2** |
| **Fmoc-L-Leucine-OH 13C6,15N1** | Fmoc-L-Leu $^{13}C_6$ $^{15}N_1$ | - | In-house synthesis, **Chapter 2** |
| ***Chemicals*** | | | |
| **(Benzotriazol-1-yloxy)tripyrrolidino-phosphonium hexafluorophosphate)** | PyBOP | 128625-52-5 | NovaBiochem, #851009 |
| **2,2′-(Ethylenedioxy)diethanethiol** | DODt | 14970-87-7 | SigmaAldrich, #465178 |
| ***N,N*-Diisopropylethylamine** | DiPEA | 7087-68-5 | VWR #84574.290 |
| **Phenol** | PhOH | 108-95-2 | SigmaAldrich, #328111 |
| **Piperidine** | - | 110-89-4 | Carlo Erba Reagents #P0663516 |
| **Trifluoroacetic acid** | TFA | 76-05-1 | Biosolve, #20233320 |
| **Triisopropylsilane** | $i$$Pr_3SiH$ | 6485-79-6 | SigmaAldrich, #233781 |
| ***Solvents*** | | | |
| **Acetonitrile (HPLC-R)** | $CH_3CN$ (AR) | 75-05-8 | VWR, #83 639 320 |
| **Acetonitrile (ULC-MS)** | $CH_3CN$ (ULC-MS) | 75-05-8 | Biosolve #1204102 |
| **Dichloromethane (AR)** | DCM | 75-09-2 | VWR, #23 366 327 |
| **Diethyl ether (AR)** | $Et_2O$ | 60-29-7 | Biosolve, #5280501 |
| **Dimethyl sulfoxide (AR)** | DMSO(AR) | 67-68-5 | Biosolve, #4470501 |
| **Formic Acid (AR)** | FA (AR) | 64-18-6 | Fisher Scientific #147932500 |
| **Formic Acid (ULC-MS)** | FA (ULC-MS) | 64-18-6 | Biosolve #6914143 |
| **Methanol** | MeOH | | VWR, #20847307 |

| | | | |
|---|---|---|---|
| ***N*-Methyl-pyrrolidone (PEPTIDE)** | NMP | 872-50-4 | VWR |
| ***n*-Pentane** | Pentane | 109-66-0 | Biosolve, #16050502 |

* Peptide building blocks are listed in Supplementary Table 8.

**Solid Phase Peptide Synthesis (SPPS)**

SPPS was performed on a Syro II Multisyntech Automated Peptide synthesizer (SYRO robot; Part Nr: S002PS002; MultiSyntech GmbH, Germany) under inert gas ($N_2$) application, using standard 9-fluorenylmethoxycarbonyl (Fmoc) based solid phase peptide chemistry on a 10 or 20 µmol scale. A fourfold excess of amino acids relative to pre-loaded Fmoc amino acid HMPA resin (between 0.17 and 0.20 mmol/g, Rapp Polymere, Germany) was used. Ubiquitin variants on resin were prepared by linear SPPS as described previously.[3] Some optimizations were made in the synthesis protocol. Optimization of the procedure led to discarding of the capping step and to the equalizing of all cycles except for the coupling cycles of isotope-labeled amino acids. Briefly, Fmoc-glycine-loaded TentaGel® R HMPA resin (Rapp Polymere, Germany, #1513) or Fmoc-Arg(Pbf)-loaded TentaGel® R HMPA resin (Rapp Polymere, Germany, #RA1502) was washed with DCM (1x 5 mL) and swelled with NMP (1x 1250 µL) for 5 minutes prior to further modifications. Fmoc-protecting groups were removed by incubating three times with 20% piperidine/NMP (v/v) for 2, 2 and 5 minutes. Resin was washed with NMP (5x 1100 µL). Fmoc-protected amino acids to-be-coupled (4 eq.) were preactivated with PyBOP (4 eq.) and DIPEA (8 eq.) in NMP. Deprotected resin was incubated twice for 25 minutes with the preactivated mixture, washed with NMP (3x 1100 µL) after the second coupling step and Fmoc removal was performed as described above. This procedure was repeated for each amino acid coupling cycle, with a total of 70, 68 or 67 cycles. Coupling cycles with isotope-labelled amino acids were extended to two times 60 minutes and only 2.8 equivalents of Fmoc-protected amino acid was used, PyBOP (4eq.) and DIPEA (8eq.) stayed unchanged. Acetylation was performed by replacing the Fmoc-protected amino acid of a standard coupling cycle with acetic anhydride (4 eq.) and no PyBOP was added in the preactivation mixture. After the two couplings cycles of acetic anhydride no deprotection cycles were performed. Details on acid-labile side chain protecting groups (PG) and coupling of Fmoc-protected (di)peptide building blocks are provided in Supplementary Table 7 and Supplementary Table 8.

**Supplementary Table 8| Fmoc protected L-amino acids and dipeptides**

| AA | Reagent | CAS# | Cat# * |
|---|---|---|---|
| **A** | Fmoc-L-Ala-OH | 35661-39-3 | 852003 |
| **R** | Fmoc-L-Arg(Pbf)-OH | 154445-77-9 | 852067 |
| **N** | Fmoc-L-Asn(Trt)-OH | 132388-59-1 | 852044 |
| **D** | Fmoc-L-Asp(OtBu)-OH | 71989-14-5 | 852005 |
| **Q** | Fmoc-L-Gln(Trt)-OH | 132327-80-1 | 852045 |
| **E** | Fmoc-L-Glu(OtBu)-OH | 71989-18-9 | 852009 |
| **G** | Fmoc-Gly-OH | 29022-11-5 | 852001 |
| **H** | Fmoc-L-His(Trt)-OH | 109425-51-6 | 852032 |
| **I** | Fmoc-L-Ile-OH | 71989-23-6 | 852010 |
| **L** | Fmoc-L-Leu-OH | 35661-60-0 | 852011 |
| **K** | Fmoc-L-Lys(Boc)-OH | 71989-26-9 | 852012 |
| **Nle** | Fmoc-L-Nle-OH | 77284-32-3 | 852014 |
| **F** | Fmoc-L-Phe-OH | 35661-40-6 | 852016 |
| **P** | Fmoc-L-Pro-OH | 71989-31-6 | 852017 |
| **S** | Fmoc-L-Ser(tBu)-OH | 71989-33-8 | 852019 |
| **T** | Fmoc-L-Thr(tBu)-OH | 71989-35-0 | 852000 |
| **Y** | Fmoc-L-Tyr(tBu)-OH | 71989-38-3 | 852020 |

| | | | |
|---|---|---|---|
| **V** | Fmoc-L-Val-OH | 68858-20-8 | 852021 |
| **AG** | Fmoc-L-Ala-(Dmb)Gly-OH | - | 852108 |
| **DG** | Fmoc-L-Asp(OtBu)- (Dmb)Gly-OH | 900152-72-9 | 852115 |
| **IT** | Fmoc-L-Ile-L-Thr($\Psi^{Me,Me}$pro)-OH | 957780-52-8 | 852193 |
| **LS** | Fmoc-L-Leu-L-Ser($\Psi^{Me,Me}$pro)-OH | 339531-50-9 | 852179 |
| **LT** | Fmoc-L-Leu-L-Thr($\Psi^{Me,Me}$pro)-OH | 955048-89-2 | 852184 |
| **ST** | Fmoc-L-Ser(tBu)-L-Thr($\Psi^{Me,Me}$pro)-OH | - | 852192 |

* Brand: Novabiochem. Supplier: Merck

**General procedure for trial cleavage**
A small amount of resin was parted from the reaction mixture and washed with DCM and $Et_2O$. The resin was air-dried, and incubated with 'fast' trial cleavage mix (TFA/$H_2O$/DODT/*i*$Pr_3SiH$; 92.5/2.5/2.5/2.5; v/v/v/v; 100 µL) and shaken for 30 minutes at 37 °C. Samples were transferred to a filter tip and filtered. The reaction mixture (filtrate) was collected in cold $Et_2O$/*n*-pentane (3/1; v/v; 1.5 mL) to precipitate the product. The suspension was centrifuged and the supernatant was decanted. The precipitate was resuspended twice in cold $Et_2O$, centrifuged and $Et_2O$ was decanted. The remaining $Et_2O$ was removed by submitting to a gentle air flow. The solid material was dissolved in DMSO (50 µL), the DMSO solution (2 µL) was diluted into 0.1% aqueous formic acid (80 µL) and reaction progress was analysed by LC-MS – System 1 – Gradient 2.

**LC-MS analysis of trial cleavages, crude reaction mixtures and purification fractions**
LC-MS analysis of crude reaction mixtures and purification fractions were performed on a Waters Alliance HT 2795 Separation Module system equipped with Waters 2996 Photodiode Array Detector (λ = 210-800 nm), Waters ACQUITY UPLC Protein BEH C4 column (300 Å, 1.7 µm, 2.1 x 50 mm) and LCT Premier Orthogonal Acceleration Time of Flight Mass Spectrometer (m/z = 100-1600) in ES+ mode (System 1). Samples were run with a 1.6 minute gradient (run time 3 min) using three mobile phases; 100% $H_2O$, 100% $CH_3CN$ and 50% $H_2O$ + 50% $CH_3CN$ + 2.5% FA (flow rate = 0.5 mL/min).

*LC-MS – System 1 – Gradient 1:*

| **Time** (min) | **100% $H_2O$** (%) | **100% $CH_3CN$** (%) | **50% $H_2O$ + 50% $CH_3CN$ + 2.5% FA**(%) |
|---|---|---|---|
| 0.00 | 94.0 | 2.0 | 4.0 |
| 0.20 | 94.0 | 2.0 | 4.0 |
| 1.80 | 0.0 | 96.0 | 4.0 |
| 2.15 | 0.0 | 96.0 | 4.0 |
| 2.20 | 94.0 | 2.0 | 4.0 |
| 3.00 | 94.0 | 2.0 | 4.0 |

*LC-MS – System 1 – Gradient 2:*

Pure products were run with a 7 minute gradient (run time 10 min) using three mobile phases: 100% $H_2O$, 100% $CH_3CN$ and 50% $H_2O$ + 50% $CH_3CN$ + 2.5% FA (flow rate = 0.5 mL/min).

| **Time** (min) | **100% $H_2O$** (%) | **100% $CH_3CN$** (%) | **50% $H_2O$ + 50% $CH_3CN$ + 2.5% FA**(%) |
|---|---|---|---|
| 0.00 | 94.0 | 2.0 | 4.0 |
| 0.50 | 94.0 | 2.0 | 4.0 |

| | | | |
|---|---|---|---|
| 7.50 | 0.0 | 96.0 | 4.0 |
| 8.00 | 0.0 | 96.0 | 4.0 |
| 8.10 | 94.0 | 2.0 | 4.0 |
| 10.00 | 94.0 | 2.0 | 4.0 |

**LC-MS analysis of purified mono- and diubiquitins**

LC-MS analysis of purified monoubiquitins and the described assay were performed on a Waters Acquitiy H-Class UPLC system equipped with a Waters ACQUITY Quaternary Solvent Manager (QSM) and Waters ACQUITY FTN AutoSampler. Separation was achieved on a Waters Acquity UPLC Protein BEH C4 column, 300Å, 1,7 uM (2.1 x 50 mm); flow rate = 0.6 mL/min , runtime = 5.55 min, column T = 60°C using 2 mobile phases: A = 0,1% formic acid in water and B = 0,1% formic acid in $CH_3CN$. Mono- and diubiquitins were separated at baseline level and eluted using a shallow gradient focused from 29% → 32% B over 2 minutes. The products were analysed by intact MS analysis (MS1) and masses were detected in a range from 550-2000 Da from 2.51 - 5.50 min and were recorded on a Waters XEVOG2 XS Q-Tof mass spectrometer equipped with an electrospray ion source in positive mode (Capillary Voltage: 0.5 kV, desolvation gas flow: 900 L/h, desolvation gas temperature: 500°C, source temperature: 130 °C, probe angle: 9.5) with a resolution of $R$ = 22,000 (System 2).

*LC-MS – System 2 – Gradient 1*: monoubiquitin gradient

| **Time** (min) | **Flowrate** (mL/min) | **100% $H_2O$** (%) **+ 0.1% FA** | **100% $CH_3CN$** (%) **+ 0.1% FA** |
|---|---|---|---|
| **0.00** | 0.6 | 98.0 | 2.0 |
| **0.20** | 0.6 | 98.0 | 2.0 |
| **0.70** | 1.0 | 98.0 | 2.0 |
| **1.80** | 1.0 | 98.0 | 2.0 |
| **2.30** | 0.6 | 98.0 | 2.0 |
| **2.50** | 0.6 | 98.0 | 2.0 |
| **2.55** | 0.6 | 71.0 | 29.0 |
| **2.83** | 0.6 | 71.0 | 29.0 |
| **4.83** | 0.6 | 68.0 | 32.0 |
| **4.90** | 0.6 | 0.0 | 100.0 |
| **5.30** | 0.6 | 0.0 | 100.0 |
| **5.35** | 0.6 | 98.0 | 2.0 |
| **5.55** | 0.6 | 98.0 | 2.0 |

In between assay runs, the column was washed using a run with the same gradient without the column flush with a 2 minutes gradient focused from 29% → 32% B (run time 2.20 min) using two mobile phases: 100% $H_2O$ + 0.1% FA and 100% $CH_3CN$ + 0.1% FA (flow rate = 0.6 mL/min) and masses were detected in a range from 550-2000 Da during the entire run time.

*LC-MS – System 2 – Gradient 2*: monoubiquitin wash run

| **Time** (min) | **Flowrate** (mL/min) | **100% $H_2O$** (%) **+ 0.1% FA** | **100% $CH_3CN$** (%) **+ 0.1% FA** |
|---|---|---|---|
| **0.00** | 0.6 | 98.0 | 2.0 |
| **0.05** | 0.6 | 71.0 | 29.0 |
| **0.33** | 0.6 | 71.0 | 29.0 |
| **2.33** | 0.6 | 68.0 | 32.0 |
| **2.40** | 0.6 | 0.0 | 100.0 |
| **2.80** | 0.6 | 0.0 | 100.0 |
| **2.85** | 0.6 | 98.0 | 2.0 |
| **3.20** | 0.6 | 98.0 | 2.0 |

Pure products were run with a 7 minute gradient (run time 10 min) using two mobile phases:
100% $H_2O$ + 0.1% FA and 100% $CH_3CN$ + 0.1% FA (flow rate = 0.5 mL/min).

*LC-MS – System 2 – Gradient 3:* Analytical run

| **Time** (min) | **Flowrate** (mL/min) | **100% $H_2O$** (%) **+ 0.1% FA** | **100% $CH_3CN$** (%) **+ 0.1% FA** |
|---|---|---|---|
| **0.00** | 0.5 | 96.0 | 4.0 |
| **0.50** | 0.5 | 96.0 | 4.0 |
| **7.50** | 0.5 | 2.0 | 98.0 |
| **8.00** | 0.5 | 2.0 | 98.0 |
| **8.10** | 0.5 | 96.0 | 4.0 |
| **10.00** | 0.5 | 96.0 | 4.0 |

NGC fractions were run with a 1.7 minute gradient (run time 3 min) using two mobile phases:
100% $H_2O$ + 0.1% FA and 100% $CH_3CN$ + 0.1% FA (flow rate = 0.6 mL/min).

*LC-MS – System 2 – Gradient 4:*

| **Time** (min) | **Flowrate** (mL/min) | **100% $H_2O$** (%) **+ 0.1% FA** | **100% $CH_3CN$** (%) **+ 0.1% FA** |
|---|---|---|---|
| **0.00** | 0.6 | 98.0 | 2.0 |
| **0.15** | 0.6 | 98.0 | 2.0 |
| **1.85** | 0.6 | 0.0 | 100.0 |
| **2.05** | 0.6 | 0.0 | 100.0 |
| **2.10** | 0.6 | 98.0 | 2.0 |
| **3.00** | 0.6 | 98.0 | 2.0 |

**RP-HPLC purification**

*System 1*. RP-HPLC purifications (max. 5 mL/run) were performed on a Shimadzu LC-20AT HPLC system equipped with a Shimadzu SPD-20A UV/Vis detector, a Shimadzu FRC-10A fraction collector and a Waters XBridge BEH C18 OBD Prep Column (130 Å, 5 µm, 10 × 150 mm) was used. Samples were run with a 15 or 23 minute gradient detailed below (run time 25 or 35 minutes) at a flowrate = 4.00 or 6.50 mL/min. Mobile phase: A = 0.05% TFA in $H_2O$ and B = 0.05% TFA in $CH_3CN$. $T$ = 40 °C.

*RP-HPLC – Gradient 1:*

| Time (min) | 0.05% TFA in $H_2O$ (%) | 0.05% TFA in $CH_3CN$ (%) | Flow rate (mL/min) |
|---|---|---|---|
| 0.00 | 95.0 | 5.0 | 4.00 |
| 6.00 | 95.0 | 5.0 | 4.00 |
| 7.00 | 90.0 | 10.0 | 6.50 |
| 10.00 | 75.0 | 25.0 | 6.50 |
| 22.00 | 50.0 | 50.0 | 6.50 |
| 22.10 | 5.0 | 95.0 | 6.50 |
| 24.00 | 5.0 | 95.0 | 6.50 |

| | | | |
|---|---|---|---|
| 24.10 | 95.0 | 5.0 | 6.50 |
| 25.00 | 95.0 | 5.0 | 6.50 |

*RP-HPLC – Gradient 2:*

| Time (min) | 0.05% TFA in $H_2O$ (%) | 0.05% TFA in $CH_3CN$ (%) | Flow rate (mL/min) |
|---|---|---|---|
| 0.00 | 95.0 | 5.0 | 4.00 |
| 6.00 | 95.0 | 5.0 | 4.00 |
| 7.00 | 90.0 | 10.0 | 6.50 |
| 10.00 | 75.0 | 25.0 | 6.50 |
| 22.00 | 65.0 | 35.0 | 6.50 |
| 30.00 | 30.0 | 70.0 | 6.50 |
| 30.10 | 5.0 | 95.0 | 6.50 |
| 32.00 | 5.0 | 95.0 | 6.50 |
| 32.10 | 95.0 | 5.0 | 6.50 |
| 35.00 | 95.0 | 5.0 | 6.50 |

Synthesis of Ac-Ub(1-74, $Nle_1$, 6x K → R, Kxx=Lys, xx V*, xx L*, xx **I***) (**4 a-g**)

**Step 1. SPPS**
See SPPS procedure with extended coupling times.
*Sequences neutron-encoded Ac-Ub1-74 $^{Met1Nle}$ (6x K → R, Kxx = Lys)*
See table Supplementary Table 1
68 cycles on Fmoc-Arg(Pbf)-loaded TentaGel® R HMPA resin (Rapp Polymere, Germany, #RA1502) on 10 µmol scale. To check the quality of the SPPS product a trial cleavage was performed.

**Step 2. Global deprotection**
Global deprotection was performed as described previously.[4] The resin-bound polypeptide Ac-Ub(1-74, $Nle_1$, 6x K → R, Kxx=Lys, xx V*, xx L*, xx **I***)(PG) **1a-g** was deprotected and detached from the resin by treatment with TFA/$H_2O$/Phenol/i$Pr_3$SiH (90.5/5/2.5/2; v/v/v/v; 2 mL) for 2.5-3.5 hours at room temperature under gentle shaking. The reaction mixture was filtered directly into ice-cold $Et_2O$/*n*-pentane (1/1; v/v; 8 mL) and the resin was washed with TFA (2x 1mL). The mixture of $Et_2O$/*n*-pentane and filtrate was centrifuged (1500 rpm, 5 min, 4 °C) and the supernatant was removed by decanting. The pellet was washed three times by resuspension in $Et_2O$ (9 mL), centrifuging (1500 rpm, 5 min, 4 °C) and removal of the supernatant. The pellet was dissolved in $H_2O$/$CH_3CN$/AcOH (75/24/1; v/v/v; 3 mL) and lyophilized. The protein was subsequently purified using RP-HPLC.

**RP-HPLC purification**
The crude monoubiquitin was properly dissolved in a minimal amount of DMSO (max. 10 vol% of the final volume) while heated carefully. The DMSO was added dropwise into $H_2O$ (10 to 20 mL). The pH was checked and should be below 7. The mixture was centrifuged (5 min @3800 rpm). The supernatant was filtered and purified by RP-HPLC.
See *RP-HPLC - Gradient 1.*

Pure fractions (checked by LC-MS) were pooled and lyophilized to obtain the product as a white powder.

Synthesis of Ub(1-74, $Nle_1$, 7x K → R, 4x V*, 6x L*, 6x **I***) (**5**)

**Step 1. SPPS**
See SPPS procedure with extended coupling times.
*Sequences neutron-encoded Ub1-74$^{Met1Nle}$(7x K→R)*
See table Supplementary Table 1
67 cycles on Fmoc-Arg(Pbf)-loaded TentaGel® R HMPA resin (Rapp Polymere, Germany, #RA1502) on 10 µmol scale. To check the quality of the SPPS product a trial cleavage was performed.

**Step 2. Global deprotection**
*vide supra* - Synthesis of Ac-Ub(1-74, $Nle_1$, 6x K → R, Kxx=Lys, xx V*, xx L*, xx **I***) (**4 a-g**)

**RP-HPLC purification**
*vide supra* - Synthesis of Ac-Ub(1-74, $Nle_1$, 6x K → R, Kxx=Lys, xx V*, xx L*, xx **I***) (**4 a-g**)

See *RP-HPLC - Gradient 1.*

Pure fractions (checked by LC-MS) were pooled and lyophilized to obtain the product as a white powder.

Synthesis of Ac-Ub(1-76, $Nle_1$, 7x K → R) (**6**)

**Step 1. SPPS**
See SPPS procedure.
*Sequences Ac-Ub1-76$^{Met1Nle}$(7x K→R)*
See table Supplementary Table 1
70 cycles on Fmoc-Gly-loaded TentaGel® R HMPA resin (Rapp Polymere, Germany, #RA1513) on 20 µmol scale. To check the quality of the SPPS product a trial cleavage was performed.

**Step 2. Global deprotection**
*vide supra* - Synthesis of Ac-Ub(1-74, $Nle_1$, 6x K → R, Kxx=Lys, xx V*, xx L*, xx **I***) (**4 a-g**)

**RP-HPLC purification**
*vide supra* - Synthesis of Ac-Ub(1-74, $Nle_1$, 6x K → R, Kxx=Lys, xx V*, xx L*, xx **I***) (**4 a-g**)

See *RP-HPLC - Gradient 1.*

Pure fractions (checked by LC-MS) were pooled and lyophilized to obtain the product as a white powder.

**Size exclusion**
The RP-HPLC products were purified by gel filtration using a Biorad NGC Chromotography system on a size exclusion S75 16/600 superdex PG-GE healthcare column with a volume bed of 120 mL and 3-70 kDa separation range using a filtered aqueous buffer containing 20 mM TRIS·HCl and 100 mM NaCl at pH 7.55 at a flowrate of 1 mL/min. The sample was prepared by dissolving the product in DMSO (250 µL), dropwise addition of this solution to MilliQ (2450 µL) and dropwise addition of 10x TRIS buffer (300 µL). The mixture was centrifuged for 5 min @3500 rpm. The fractions were analysed by SDS-PAGE and LC-MS and pure fractions were pooled. The products were obtained as colorless solutions containing 20 mM TRIS·HCl and 100 mM NaCl buffer at pH 7.55. LC-MS analysis (LC-MS – System 2 – Gradient 4) was done to check the purity. Pure fractions were combined and concentrated using 3 kDa cut-off spin filters.

**Concentration determination**
To determine the concentration of the solution (and the yield), this solution was together with a concentration series of mono-ubiquitin (0.5 µg, 1 µg, 2 µg, 3 µg per lane), resolved by SDS-PAGE, stained with InstantBlue™ Staining and scanned. The concentration of the solution was determined by quantification of the bands using a GE Healthcare Amersham Imager 600 with ImageQuant TL 8.1 GE Healthcare lifesciences software.

| Protein | Stock concentration (mg/mL) | Stock concentration (µM) | Amount |
|---|---|---|---|
| **Neutron-encoded K6 monoUb (4a)** | 1.27 | 146.9 | 1030 µL |
| **Neutron-encoded K11 monoUb (4b)** | 2.01 | 232.2 | 1120 µL |
| **Neutron-encoded K27 monoUb (4c)** | 1.87 | 215.7 | 1000 µL |
| **Neutron-encoded K29 monoUb (4d)** | 1.88 | 216.6 | 1100 µL |
| **Neutron-encoded K33 monoUb (4e)** | 3.32 | 381.8 | 900 µL |
| **Neutron-encoded K48 monoUb (4f)** | 3.55 | 407.6 | 1000 µL |
| **Neutron-encoded K63 monoUb (4g)** | 0.88 | 100.9 | 1700 µL |
| **Neutron-encoded M1 monoUb (5)** | 1.30 | 148.8 | 1400 µL |
| **Donor monoUb (6)** | 1.16 | 132.04 | 1400 µL |

Ac-Ub(1-74, $Nle_1$, 6x K → R, Kxx=Lys, xx V*, xx L*, xx **I***) (**4 a-g**)
The products were obtained as white solids. LC-MS analysis using System 2 – Gradient 3.
Yields:
Ac-Ub(1-74, $Nle_1$, 6x K→R, K6=Lys) **4a** = 1.31 mg, 0.15 µmol, 1.5%. LC-MS: $R_t$ 3.17 min: MS ES+ (amu) calculated: 8642,85Da[M]; found: 8646 Da.
Ac-Ub(1-74, $Nle_1$, 6x K→R, K11=Lys 1x V*, 1x L*) **4b** = 2.25 mg, 0.26 µmol, 2.6%. LC-MS: $R_t$ 3.17 min: MS ES+ (amu) calculated: 8655,75Da[M]; found 8657 Da.
Ac-Ub(1-74, $Nle_1$, 6x K→R, K27=Lys 3x V*, 1x **I***) **4c** 1.20 mg, 0.14 µmol, 1.4%. LC-MS: $R_t$ 3.17 min: MS ES+ (amu) calculated: 8667.66 Da[M]; found 8669 Da.
Ac-Ub(1-74, $Nle_1$, 6x K→R, K29=Lys 3x V*, 2x L*, 1x **I***) **4d** = 2.07 mg, 0.24 µmol, 2.4%. LC-MS: $R_t$ 3.17 min: MS ES+ (amu) calculated: 8681.55 Da[M]; found 8682 Da.
Ac-Ub(1-74, $Nle_1$, 6x K→R, K33=Lys 3x V*, 2x L*, 3x **I***) **4e** = 2.99 mg, 0.34 µmol, 3.4%. LC-MS: $R_t$ 3.17 min: MS ES+ (amu) calculated: 8695.44 Da[M]; found 8695 Da.
Ac-Ub(1-74, $Nle_1$, 6x K→R, K48=Lys 3x V*, 4x L*, 3x **I***) **4f** = 3.55 mg, 0.41 µmol, 4.1%. LC-MS: $R_t$ 3.17 min: MS ES+ (amu) calculated: 8709.33Da[M]; found 8711 Da.
Ac-Ub(1-74, $Nle_1$, 6x K→R, K63=Lys 3x V*, 4x L*, 5x **I***) **4g** = 1.50mg, 0.17 µmol, 1.7%. LC-MS: $R_t$ 3.17 min: MS ES+ (amu) calculated: 8723.22Da[M]; found 8723 Da.

Ub(1-74, $Nle_1$, 7x K → R, 4x V*, 6x L*, 6x **I***) (**5**)
The products were obtained as white solids. LC-MS analysis using System 2 – Gradient 3.
Yield:
Ub(1-74, $Nle_1$, 7x K → R, 4x V*, 6x L*, 6x **I***) **5 =** 1.82 mg, 0.21 µmol, 2.1%. LC-MS: $R_t$ 3.03 min: MS ES+ (amu) calculated: 8736.04 [M]; found 8736 Da.

Ac-Ub(1-76, $Nle_1$, 7x K → R) (**6**)
The products were obtained as white solids. LC-MS analysis using System 2 – Gradient 3.
Yield:
Ac-Ub(1-76, $Nle_1$, 7x K → R) **6 =** 1.6 mg, 0.18 µmol. LC-MS: $R_t$ 3.17 min: MS ES+ (amu) calculated: 8784.94 [M]; found 8788 Da.

Synthesis of Ub(1-74, $Nle_1$)
Synthesis of Ub(1-74, $Nle_1$) was performed as described previously (see **Chapter 2**).
In short, Ub(1-74, $Nle_1$) was synthesized using SPPS (see SPPS procedure).
*Sequence Ub1-74$^{Met1Nle}$*

(Nle)QIFVKTLTG KTITLEVEPS DTIENVKAKI QDKEGIPPDQ QRLIFAGKQL EDGRTLSDYN IQKESTLHLV LRLR
67 cycles on Fmoc-Arg(Pbf)-loaded TentaGel® R HMPA resin (Rapp Polymere, Germany, #RA1502) 25 µmol scale
Afterwards, Ub(1-74, $Nle_1$) was detached from the resin and globally deprotected. The obtained solid crude material was lyophilized and purified using RP-HPLC. Pure fractions (>95%, checked by LC-MS) were pooled and lyophilized to obtain the product as a white powder.

Ub(1-74, $Nle_1$)

The product was obtained as white solid. LC-MS analysis using LC-MS – System 2 – Gradient 3. Yield:

Ub(1-74, $Nle_1$) = 32.85 mg, 3.89 µmol, 31.1%. LC-MS: $R_t$ 3.13 min: MS ES+ (amu) calculated: 8432.7 Da[M]; found 8433 Da.

The products were purified by gel filtration using a Biorad NGC Chromotography system on a size exclusion S75 16/600 superdex PG-GE healthcare column. See **Size Exclusion** (above). The product was obtained as colorless solution containing 50 mM TRIS·HCl and 20 mM NaCl buffer at pH 7.55. LC-MS analysis (LC-MS – System 2 – Gradient 4) was done to check the purity. Pure fractions were combined and concentrated using 3 kDa cut-off spin filters. Yielding a stock concentration of 2.5 mg/mL or 296.5 µM.

Synthesis of non-hydrolysable clicked Lys48 diubiquitin

The synthesis of non-hydrolysable clicked Lys48 diubiquitin was performed as described previously (see **Chapter 2**).
In short, Ub1-75$^{Met1Nle}$ and Ub1-75$^{Met1Nle}$ (K48 = L-azido-ornithine) were synthesized using SPPS (see SPPS procedure).
*Sequence Ub1-75$^{Met1Nle}$*
(Nle)QIFVKTLTG KTITLEVEPS DTIENVKAKI QDKEGIPPDQ QRLIFAGKQL EDGRTLSDYN IQKESTLHLV LRLRG
Sequence Ub1-75$^{Met1Nle}$ (K48 = L-azido-ornithine)
(Nle)QIFVKTLTG KTITLEVEPS DTIENVKAKI QDKEGIPPDQ QRLIFAG(L-azido-ornithine)QL EDGRTLSDYN IQKESTLHLV LRLRG
68 cycles on Fmoc-Gly-loaded TentaGel® R trityl resin (Rapp Polymere, Germany, #RA RA1213) 25 µmol scale

Afterwards, Ub1-75$^{Met1Nle}$ (K48 = L-azido-ornithine) was detached from the resin and globally deprotected. The obtained solid crude material was lyophilized and purified using RP-HPLC (*RP-HPLC - Gradient 1*). Pure fractions (>95%, checked by LC-MS) were pooled and lyophilized to obtain the product as a white powder.

Ub(1-75, $Nle_1$)-PA was prepared as described previously.[5] Briefly, after SPPS and release from the resin using HFIP/DCM (1/4; v/v) the protected polypeptide Ub(1-75, $Nle_1$) (25 µmol) was dissolved in DCM (1 mL/ 5 µmol), and PyBOP (5 eq., 125 µmol), triethylamine (5 eq., 125 µmol) and propargylamine (10 eq., 250 µmol) were added to the solution. The reaction was stirred for 16 hours at RT. The reaction mixture was concentrated and the polypeptide was globally deprotected. The obtained solid crude material was lyophilized and purified using preparative RP-HPLC (*RP-HPLC - Gradient 1*). Pure fractions (> 95%, checked by LC-MS) were pooled and lyophilized to obtain the product as a white powder.

Ub1-75$^{Met1Nle}$ (K48 = L-azido-ornithine) and Ub(1-75, $Nle_1$)-PA were clicked together as described previously.[6] Briefly, the CuAAC reactions were performed under denaturing conditions in 8 M Urea, 100 mM phosphate buffer pH 7. Ub(1-75, $Nle_1$)-PA (11.5 mg) was dissolved in warm DMSO (100 µL) and Ub(1-75, $Nle_1$, L-azido-ornithine$_{48}$) (10.25 mg) was dissolved in warm DMSO (100 µL). Both DMSO solutions were added to an aqueous buffer containing 8 M Urea and 100 mM phosphate pH 7 (2 mL). To the solution 210 µL of catalyst solution containing 25 mg/mL $CuSO_4·5H_2O$ in $H_2O$, 120 mg/mL sodium ascorbate in $H_2O$ and 52 mg/mL TBTA-analogue[7] in $CH_3CN$ (1/1/1; *v/v/v*) was added. The reaction was gently shaken at room temperature. Extra catalyst solution was added after 1h (90 µL) and fresh catalyst solution was added after 16h (210 µL). After reaction was finished, as judged by LC-MS (~ 24 hour), the reaction was quenched by the addition of 34 µL of 0.5 M EDTA, pH 7.0. The crude material was subsequently purified using preparative RP-HPLC (*RP-HPLC - Gradient 2*). Pure fractions (checked by *LC-MS – System 1 – Gradient 1*) were pooled and lyophilized to obtain the product as a white powder.

The products were purified by gel filtration using a Biorad NGC Chromotography system on a size exclusion S75 16/600 superdex PG-GE healthcare column. See **Size Exclusion** (above). The product was obtained as colorless solution containing 50 mM TRIS·HCl and 20 mM NaCl buffer at pH 7.55. LC-MS analysis (*LC-MS – System 2 – Gradient 4*) was done to check the purity. Pure fractions were combined and concentrated using 3 kDa cut-off spin filters. Yielding a stock concentration of 1.52 mg/mL or 89.25 µM.

#### Materials and Methods; Biochemistry

##### Recombinant protein expression and purification.

*Protein expression constructs.* The human full length TRIM25 (1-630) was cloned from Addgene plasmid #12449 into our modified bacterial pGEX6p-1 expression vector with a 3C-cleavable N-terminal GST tag. Expression vectors for the human E2 enzymes, pET28a-LIC_Uev1a (UBE2V1) (Addgene plasmid # 25619) and pET-SUMO_Ubc13 (UBE2N) (Addgene plasmid # 51131) were kind gifts from Cheryl Arrowsmith and Cynthia Wolberger respectively. Expression vectors for the E3 enzymes, pMCSG17-NleL (Addgene plasmid #66716), pOPINS-AREL1 (Addgene plasmid #66710) and pOPINS-UBEC3 (Addgene plasmid #66711) were all kind gifts from David Komander.

*Protein expression.* Overexpression of all proteins was done in *Escherichia coli* BL21-CodonPlus (DE3)-RIL strain. For TRIM25, cells were grown in 2xYT media supplemented with 50 µM $ZnCl_2$ in a shaker incubator. Growth was allowed at 37°C until $OD_{600}$ was nearly 1.0. The culture was then cooled to 18°C, expression was induced with 0.3 mM Isopropyl–β-D thiogalactopyranoside (IPTG) and left to incubate at 16 °C overnight. For UBE2N and UBE2V1, LB media was used and growth was allowed at 37°C until $OD_{600}$ reached 1.2. Induction was done with 0.3 mM IPTG at 25°C for 5 hrs for UBE2V1 and 30°C for 3 hrs for UBE2N with shaking at 200 rpm. For NleL and UBE3C, LB media was used and for AREL1, 2xYT media was used. Growth was allowed at 37°C until $OD_{600}$ reached 0.6 (NleL & UBE3C) and 1.6 (AREL1). Cultures were cooled to 18°C prior to overnight induction with 400 µM IPTG.
TRIM25 expressing cells were harvested and resuspended in lysis buffer (50 mM HEPES pH 7.5, 500 mM NaCl, 10% glycerol). Just prior to lysis, the suspension was supplemented with 1 mM fenylmethylsulfonylfluoride (PMSF) and benzamidine. UBE2N and UBE2V1 expressing cells were harvested and resuspended in lysis buffer (50 mM HEPES pH 7.5, 200 mM NaCl) having 1 mM PMSF and benzamidine. For all of them, lysis was done using Fisherbrand Q125 sonicator for 3 mins with 15-/45-s on/off cycles. The lysate was centrifuged at 14,000 x g for 40 mins at 4°C.
NleL expressing cells were harvested and resuspended in GST buffer (50 mM HEPES pH 7.5, 250 mM NaCl, 1 mM EDTA, 1 mM DTT). Lysis was done using Fisherbrand Q125 sonicator for 5 mins with 15-/45-s on/off cycles. The lysate was centrifuged at 14,000 x g for 40 mins at 4°C.
UBE3C expressing cells were harvested and resuspended in lysis buffer (20 mM TRIS pH 8.5, 300 mM NaCl, 50 mM imidazole). 1 mM PMSF (1000x in isopropanol) and 2 mM TCEP were added to the buffer. Lysis was done using Fisherbrand Q125 sonicator for 5 mins with 15-/45-s on/off cycles. The lysate was centrifuged at 14,000 x g for 40 mins at 4°C.
AREL1 expressing cells were harvested and resuspended in lysis buffer (20 mM TRIS pH 8.5, 300 mM NaCl, 50 mM imidazole). No PMSF or TCEP was added to the buffer. Lysis was done using Fisherbrand Q125 sonicator for 5 mins with 15-/45-s on/off cycles. The lysate was centrifuged at 24,000 x g for 45 mins at 4°C.

*Purification.* To purify TRIM25 and NleL, supernatant was allowed to flow through Glutathion Sepharose 4B beads (GE Healthcare). Washing was done using Wash buffer (50 mM HEPES pH 7.5, 250 mM NaCl, (1 mM EDTA for NleL), and 1 mM DTT) twice. Proteins were eluted with Wash buffer supplemented with 20 mM (TRIM25) or 25 mM (NleL) GSH. Eluted TRIM25 fractions were subjected to 3C proteolytic cleavage under dialysis against 20 mM HEPES pH 7.0, 100 mM NaCl, 1 mM DTT, 5% glycerol overnight at 4°C. The

dialysed sample was loaded on a HiTrap SP FF ion exchange column (Cytiva) using IEX buffer A (20 mM HEPES pH 7.0, 1 mM DTT, 5% glycerol). TRIM25 protein was eluted using a gradient with buffer B (20 mM HEPES pH 7.0, 1M NaCl, 1 mM DTT, 5% glycerol) and protein-containing fractions were pooled and concentrated using a 50 kDa MWCO Amicon Ultra spin concentrator, before being aliquoted and flash-frozen for storage at -80°C.
Eluted NleL fractions were subjected to TEV proteolytic cleavage under dialysis against 20 mM HEPES pH 7.5, 50 mM NaCl, 1 mM DTT overnight at 4°C. The dialysed sample was loaded on a HiTrap monoQ column (GE Healthcare). Protein was eluted using a salt gradient (20 mM HEPES pH7.5, 1 mM DTT, 50 to 1000 mM NaCl) and protein-containing fractions were pooled and concentrated using a 50 kDa MWCO Amicon Ultra spin concentrator. Concentrated fractions were purified over a Superdex75 10/30 gel filtration column equilibartad in 20 mM HEPES pH 7.5, 50 mM NaCl, 1 mM DTT. NleL was concentrated using a 50 kDa MWCO Amicon Ultra spin concentrator and aliquoted before being flash frozen for storage at -80°C.

Purification of UBE2N and UBE2V1 was done by Ni-NTA affinity chromatography. Supernatant was applied to nickel-charged chelating sepharose beads (GE Healthcare). Beads were washed with buffer A (20 mM HEPES pH7.5, 350 mM NaCl, 20 mM Imidazole, 2 mM DTT) before elution with the same buffer supplemented with 200 mM Imidazole. Following this, for UBE2N, 6xHis-SUMO tag was cleaved using SENP2CD (Sentrin/SUMO- specific protease) along with overnight dialysis against 20 mM HEPES pH 7.5, 100 mM NaCl. Similar dialysis conditions were used for UBE2V1. Next day, the proteins were further purified on a Superdex 75 10/300 GL column (GE Healthcare) equilibrated in 20 mM HEPES pH7.5, 100 mM NaCl and 2 mM DTT. Protein-containing fractions were concentrated using a 10 kDa MWCO Amicon Ultra spin filter and aliquots were flash-frozen and stored at -80°C.
Purification of UBE3C and AREL1 was done by Ni-NTA affinity chromatography. Supernatant was applied to nickel-charged chelating sepharose beads (GE Healthcare). Beads were washed with buffer A (20 mM TRIS pH8.5, 350 mM NaCl, 50 mM Imidazole) before elution with the same buffer supplemented with 200 mM imidazole. Following this, for UBE3C, 6xHis-SUMO tag was cleaved using SENP2CD (Sentrin/SUMO-specific protease) along with overnight dialysis against 20 mM TRIS pH 8.5, 150 mM NaCl, 2mM DTT. Next day, the protein was further purified on a Superdex 75 10/300 GL column (GE Healthcare) equilibrated in 20 mM TRIS pH8.5, 150 mM NaCl and 2 mM DTT. Protein-containing fractions were concentrated using a 10 kDa MWCO Amicon Ultra spin filter and aliquots were flash-frozen and stored at -80°C. For AREL1, 6xHis-SUMO tag was cleaved using SENP2CD (Sentrin/SUMO- specific protease) along with overnight dialysis against 20 mM TRIS pH 8.5, 150 mM NaCl, 2mM DTT. The dialysed sample was loaded on a HiTrap monoQ column (GE Healthcare). Protein was eluted using a salt gradient (20 mM TRIS pH 8.5, 1 mM DTT, 50 to 1000 mM NaCl) and protein-containing fractions were pooled and concentrated. The protein was further purified on a Superdex 75 10/300 GL column (GE Healthcare) equilibrated in 20 mM TRIS pH 8.5, 150 mM NaCl and 2 mM DTT. Protein-containing fractions were concentrated using a 10 kDa MWCO Amicon Ultra spin filter and aliquots were flash-frozen and stored at -80°C.

**General method SDS-PAGE analysis**
After indicated reaction time, the reaction was quenched by addition of 3x reducing sample buffer (SB) (containing 900 µL 4x LDS sample buffer (NuPAGE, Invitrogen) diluted with 210 µL water and 90 µL β-mercaptoethanol) and heated to 95°C for 5 min. (denaturing conditions). Samples were loaded on precast 4-12% or 12% NuPAGE® Novex® Bis-Tris Mini Gels (Invitrogen) and resolved by SDS-PAGE gel electrophoresis using MES running buffer (NuPAGE MES SDS running buffer 20X, Novex by Life Technologies). Reference protein standard/ladder: SeeBlue™ Plus2 Pre-stained Protein Standard (Invitrogen, cat# LC5925). Proteins were visualized by InstantBlue™ (Expedeon Protein Solutions, #ISB1L), and stained gels were scanned using a GE Healthcare Amersham Imager 600.

**Characterization of all eight neutron-encoded monoUbs, donor monoUb, K48 click diUb and $Ub_{1\text{-}74}$ by SDSPAGE analysis**
All eight neutron-encoded monoUbs, donor monoUb, K48 click diUb and $Ub_{1\text{-}74}$ stock solutions were diluted to ~3.5 µM in a buffer containing 20 mM Tris·HCl, 100 mM NaCl, pH 7.55. To 10 µL of these solutions 5 µL 3x SB was added, samples were boiled and loaded on gel (7.5 µL/lane), separated by gel electrophoresis and stained with InstantBlue staining. (**Fig. 2c** and **Supplementary Figure 1**).

**SDS-PAGE analysis of E1-E2(-E3) ubiquitin chain conjugation**

All neutron-encoded monoUbs, donor monoUb and Ub WT were not diluted. UBE1 was diluted in a buffer containing 20 mM Tris·HCl, 100 mM NaCl, pH 7.55, 5 mM $MgCl_2$, 5 mM TCEP (1 µM 10x final conc.). E2/E3 fusion ligases Ube2G2-gf7826 and UBE2S-IsoT were diluted in a buffer containing 20 mM Tris·HCl, 100 mM NaCl, pH 7.55, 5 mM $MgCl_2$, 5 mM TCEP (20 µM 10x final conc.). Subsequently, 5 µL UBE1, 5 µL E2 or E2-E3 fusion, *x* µL monoUb substrate and 40-*x* µL buffer (20 mM Tris·HCl, 100 mM NaCl, pH 7.55, 5 mM $MgCl_2$, 5 mM TCEP) were mixed to yield a solution containing 100 nM UBE1, 2 µM E2 or E2-E3 fusion and 40 or 80 µM monoUb substrate. (Ub WT 80 µM, donor Ub 40 µM, acceptor monoUb 8 x 5 µM or 40 µM). 0.5 µL 0.5 M ATP was added to start the conjugation reaction. After 30 min, 0.166 µL 0.5 M ATP was added to the reaction mixture. Samples for timepoint 0 min. were taken from the reaction mixture before the addition of ATP and quenched with 3x SB (5 µL). Samples for other timepoints (10 min., 30 min. and 60 min.) were taken from the reaction mixture and quenched with 3x SB (5 µL) and analysed according to the general method for SDS-PAGE analysis (**Fig. 3**, **Supplementary Figure 3** and **5**).

***In vitro* conjugation assays with mass spectrometry read-out**

The assays were performed in a 1.5 mL Eppendorf tube at 37°C in a buffer containing 20 mM Tris·HCl, 100 mM NaCl, pH 7.55 and 5.0 mM final concentration of TCEP and $MgCl_2$. All eight neutron-encoded monoubiquitins were mixed in an equimolar amount (8x ~7.5 µM; 1.5x final concentration) together with the donor monoubiquitin (60 µM; 1.5x final concentration). Neutron-encoded K6 (16 µL), K11 (9 µL), K27 (11 µL), K29 (11 µL), K33 (6 µL), K48 (6µL), K63 (13 µL), M1 (8 µL), donor Ub (139.2 µL) stocks were mixed and diluted with buffer (20 mM Tris·HCl, 100 mM NaCl, pH 7.55; 80.8 µL). Recombinant purified conjugation enzymes were obtained from commercial sources, received as a gift, or expressed and purified according to reported procedures (details provided in **Supplementary Table 2** and **3**). Recombinant purified conjugation enzymes were diluted to 10x final concentration (1 µM for UBE1, 20, 60 or 100 µM for E2 enzymes and 20, 40, 120 or 200 µM for E3 ligases) in a buffer containing 20 mM Tris·HCl, 100 mM NaCl, pH 7.55 and **50** mM TCEP and $MgCl_2$. The monoUb mixture (6.33 µL) was added to the Eppendorf tube. Subsequently, the enzymes (1 µL E1, 1 µL E2 and 1 µL E3, 10x final concentration) were added to the monoUb mixture. The mixture was diluted with 0.7 µL buffer (20 mM Tris·HCl, 100 mM NaCl, pH 7.55 and **50** mM TCEP and $MgCl_2$) or 1.7 µL when no E3 ligase was present. The reaction was started by the addition of 0.2 µL 0.5 M ATP. The reaction mixtures were incubated at 37°C for 180 min or 24 hours. After 30 and 60 minutes fresh ATP (0.2 µL 0.5 M) was added to the solution. Before the addition of ATP, 1 µL of reaction mixture was taken for timepoint 0 minutes. After 10, 30, 60, 180 min (and 24 hours) a sample was taken from the reaction mixture for analysis.

*Sample preparation.* After indicated incubation time, 1 µL of the reaction mixture was taken. The reaction was quenched by acidification of the mixture and internal standards for MS analysis were directly added. Therefore, 1 µL of the reaction mixture was quenched, spiked and diluted with a mixture containing 0.2 µL 10% TFA in MQ, 12.8 µL 0.1% FA in MQ and 10 µL of the internal standard solution (1 µL of 5.0 µM internal standard diluted with 9 µL of 0.1% FA in MQ). Samples were collected in a 96-well plate and measured by LC-MS analysis. 4 µL of the sample was injected onto the column.

*Data acquisition.* The samples were separated by an Acquity H-class UPLC system using a BEH C4 column (300Å, 1.7 µM (2.1 x 50 mm)), column T = 60°C. For the first 2.5 min, the flow was diverted from the detector to flush the column with 2% ACN in $H_2O$ and 0.1% FA at 1 mL/min to elute most of the buffer components and salt. After 2.5 min. proteins were eluted using a shallow gradient that ranged from 29% up to 32% ACN in $H_2O$ with 0.1% FA over 2 min using a flow rate of 0.6 mL/min, which was able to separate monoUb and diUb products (baseline level). Products were analysed by intact MS analysis on a XEVO G2 XS Q-TOF in reflector positive ion mode with a resolution of $R$ = 22,000 using positive electrospray ionization (ESI) (Cap. V. = 0.5 kV) and a detection range of $m/z$ 550-2000. The system check of the detector voltage, lock mass accuracy (of LeuEnk) and calibration using NaI solution were performed daily prior to analysis. Lock mass correction was applied during each analysis to correct for possible mass shifts during the course of the assay.

*Data analysis.* Proteowizard 3.0.20274 was used to convert raw data files to the mzXML file format.[8] mzXML files were further processed using LaCyTools version 22.04.29. The alignment was performed using theoretical $m/z$ values of various charge states from the internal standard and the expected elution time (**Supplementary Table 4**). The extraction parameters are specified in **Supplementary Table 5**. Atom compositions were determined of all eight diUb molecules, all nine monoUbs, $Ub_{1-74}$ as well as the non-hydrolysable clicked K48 diUb and listed as analytes (**Supplementary Table 6**). From all analytes, their retention time was predicted and the charge states that should be taken along during quantification are specified (**Supplementary Table 6**). The LaCyTools output file (Summary.txt) was further processed in Microsoft Excel, where boundaries for quality control parameters (Mass Accuracy < 15 ppm; IPQ < 0.25; S/N > 9) were set and the areas of all *m/z* peaks, within the quality control boundaries, from the same analyte were summed. The absolute area for each analyte was normalized at each timepoint using the internal standard, nonhydrolyzable clicked Lys48 diUb. The concentration of diUb present at each timepoint was calculated using the theoretical concentration of present nonhydrolyzable clicked Lys48 diUb in the analysed mixture. The concentration of diUb was plotted against time using GraphPadPrism 9.3.0. For the monoUb signals, the total absolute area under the curve was normalized at each timepoint using the internal standard $Ub_{1-74}$. The concentration of monoUb present at each timepoint was calculated using the theoretical concentration of present $Ub_{1-74}$ in the analysed mixture. Concentration monoUb was plotted against time using GraphPadPrism 9.3.0.

**Code availability.** LaCyTools is freely available for download at https://github.com/Tarskin/LaCyTools.

#### References


1. Mulder, M. P. C. *et al.* A cascading activity-based probe sequentially targets E1-E2-E3 ubiquitin enzymes. *Nat. Chem. Biol.* **12**, 523–530 (2016).
2. Liwocha, J. *et al.* Linkage-specific ubiquitin chain formation depends on a lysine hydrocarbon ruler. *Nat. Chem. Biol.* (2020) doi:10.1038/s41589-020-00696-0.
3. El Oualid, F. *et al.* Chemical synthesis of ubiquitin, ubiquitin-based probes, and diubiquitin. *Angew. Chemie - Int. Ed.* **49**, 10149–10153 (2010).
4. Geurink, P. P. *et al.* Development of Diubiquitin-Based FRET Probes To Quantify Ubiquitin Linkage Specificity of Deubiquitinating Enzymes. *ChemBioChem* **17**, 816–820 (2016).
5. Ekkebus, R. *et al.* On Terminal Alkynes That Can React with Active-Site Cysteine Nucleophiles in Proteases. *J. Am. Chem. Soc.* **135**, 2867–2870 (2013).
6. Flierman, D. *et al.* Non-hydrolyzable Diubiquitin Probes Reveal Linkage-Specific Reactivity of Deubiquitylating Enzymes Mediated by S2 Pockets. *Cell Chem. Biol.* **23**, 472–482 (2016).
7. Zhou, Z. & Fahrni, C. J. A Fluorogenic Probe for the Copper(I)-Catalyzed Azide−Alkyne Ligation Reaction: Modulation of the Fluorescence Emission via 3 (n,π*) − (π,π*) Inversion. *J. Am. Chem. Soc.* **126**, 8862–8863 (2004).
8. Kessner, D., Chambers, M., Burke, R., Agus, D. & Mallick, P. ProteoWizard: open source software for rapid proteomics tools development. *Bioinformatics* **24**, 2534–2536 (2008).

### *UBE2L3 + NleL*

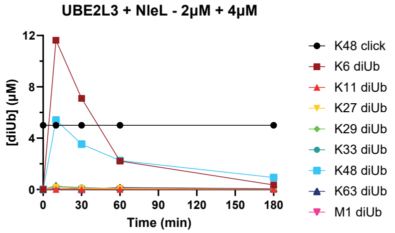
UBE2L3 + NleL - 2µM + 4µM
[diUb] (µM)
Time (min)
0
4
8
12
30
60
90
120
150
180
K48 click
K6 diUb
K11 diUb
K27 diUb
K29 diUb
K33 diUb
K48 diUb
K63 diUb
M1 diUb

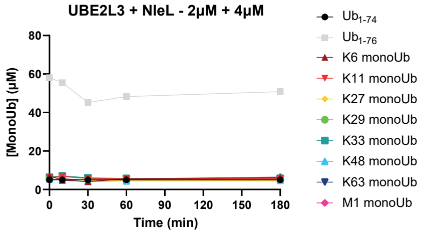
UBE2L3 + NleL - 2µM + 4µM
[MonoUb] (µM)
Time (min)
0
20
40
60
80
30
90
120
150
180
Ub1-74
Ub1-76
K6 monoUb
K11 monoUb
K27 monoUb
K29 monoUb
K33 monoUb
K48 monoUb
K63 monoUb
M1 monoUb

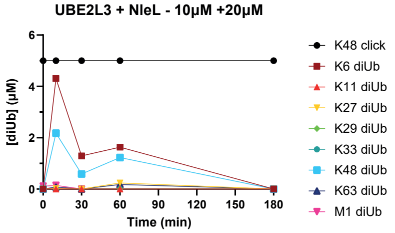
UBE2L3 + NleL - 10µM +20µM
[diUb] (µM)
Time (min)
0
2
4
6
30
60
90
120
150
180
K48 click
K6 diUb
K11 diUb
K27 diUb
K29 diUb
K33 diUb
K48 diUb
K63 diUb
M1 diUb

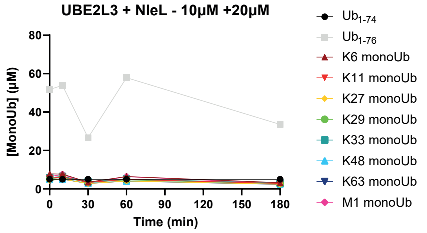
UBE2L3 + NleL - 10µM +20µM
[MonoUb] (µM)
Time (min)
0
20
40
60
80
30
90
120
150
180
Ub1-74
Ub1-76
K6 monoUb
K11 monoUb
K27 monoUb
K29 monoUb
K33 monoUb
K48 monoUb
K63 monoUb
M1 monoUb

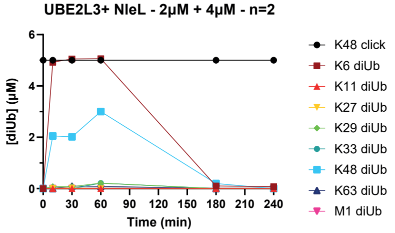
UBE2L3+ NleL - 2µM + 4µM - n=2
[diUb] (µM)
Time (min)
0
2
4
6
30
60
90
120
150
180
210
240
K48 click
K6 diUb
K11 diUb
K27 diUb
K29 diUb
K33 diUb
K48 diUb
K63 diUb
M1 diUb

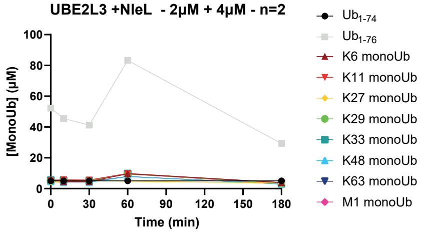
UBE2L3 +NleL - 2µM + 4µM - n=2
[MonoUb] (µM)
Time (min)
0
20
40
60
80
100
30
90
120
150
180
Ub1-74
Ub1-76
K6 monoUb
K11 monoUb
K27 monoUb
K29 monoUb
K33 monoUb
K48 monoUb
K63 monoUb
M1 monoUb

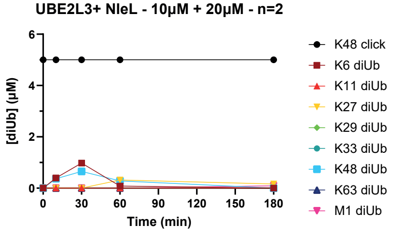
UBE2L3+ NleL - 10µM + 20µM - n=2
[diUb] (µM)
Time (min)
0
2
4
6
30
60
90
120
150
180
K48 click
K6 diUb
K11 diUb
K27 diUb
K29 diUb
K33 diUb
K48 diUb
K63 diUb
M1 diUb

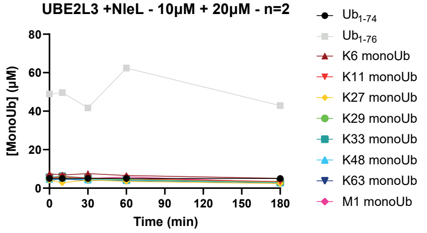
UBE2L3 +NleL - 10µM + 20µM - n=2
[MonoUb] (µM)
Time (min)
0
20
40
60
80
30
90
120
150
180
Ub1-74
Ub1-76
K6 monoUb
K11 monoUb
K27 monoUb
K29 monoUb
K33 monoUb
K48 monoUb
K63 monoUb
M1 monoUb

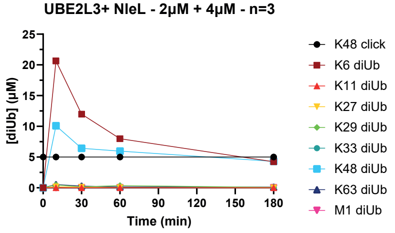
UBE2L3+ NleL - 2µM + 4µM - n=3
[diUb] (µM)
Time (min)
0
5
10
15
20
25
30
60
90
120
150
180
K48 click
K6 diUb
K11 diUb
K27 diUb
K29 diUb
K33 diUb
K48 diUb
K63 diUb
M1 diUb

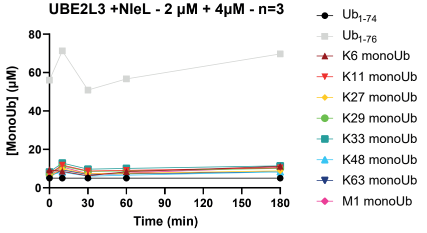
UBE2L3 +NleL - 2 µM + 4µM - n=3
[MonoUb] (µM)
Time (min)
0
20
40
60
80
30
90
120
150
180
Ub1-74
Ub1-76
K6 monoUb
K11 monoUb
K27 monoUb
K29 monoUb
K33 monoUb
K48 monoUb
K63 monoUb
M1 monoUb

#### *UBE2L3 + NleL*

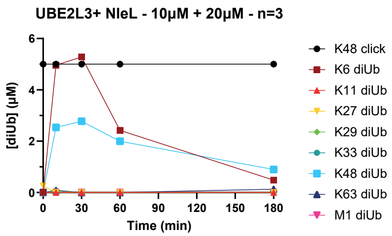


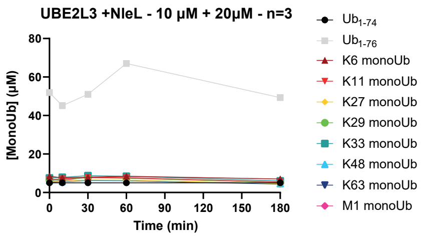


#### *UBE2S-IsoT*

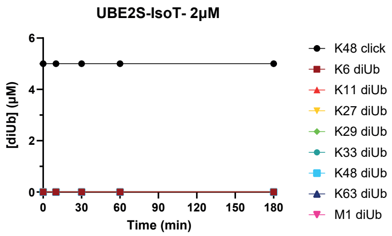


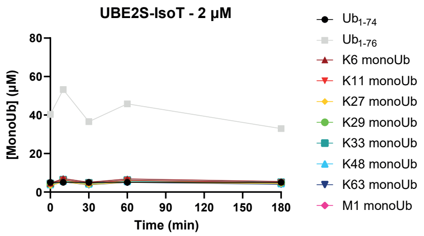


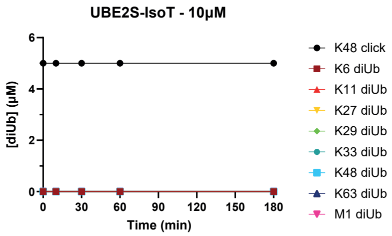


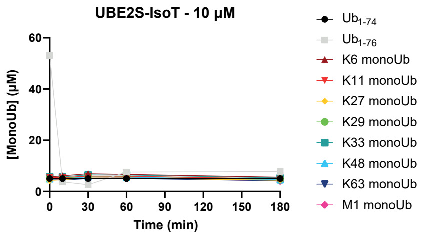


#### *UBE2L3 + UBE3C*

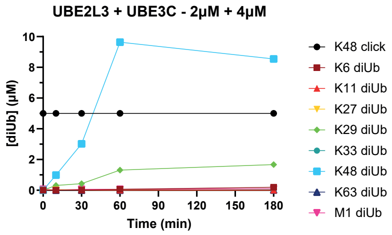


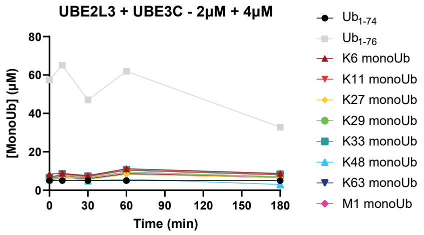


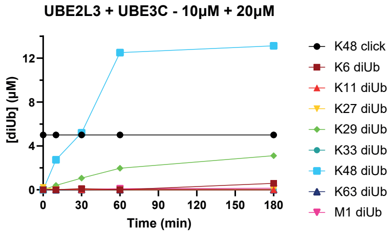


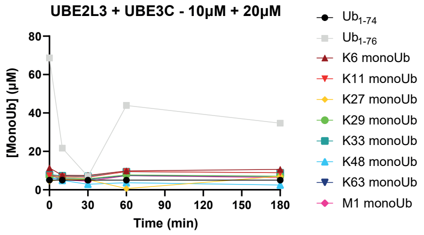

### UBE2L3 + AREL1

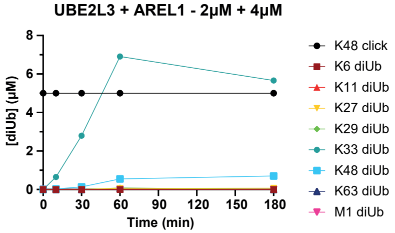


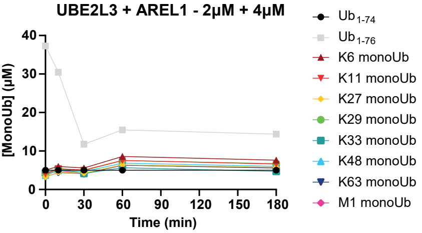


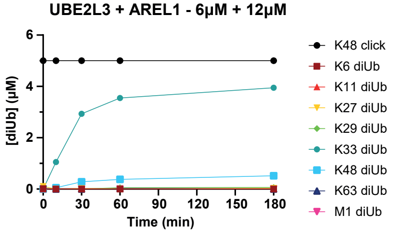


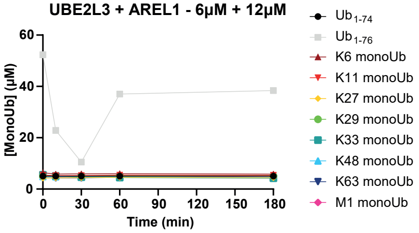


### UBE2R1

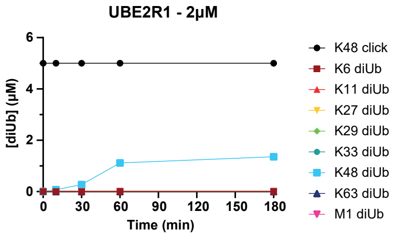


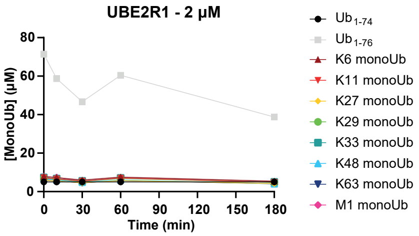


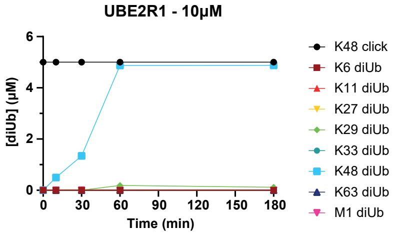


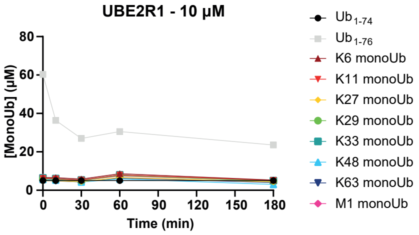


### UBE2R2

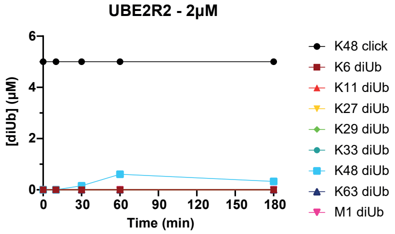


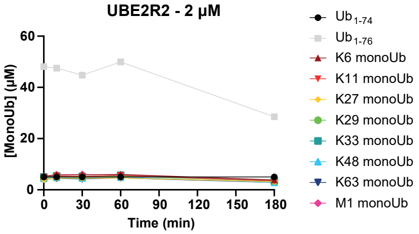

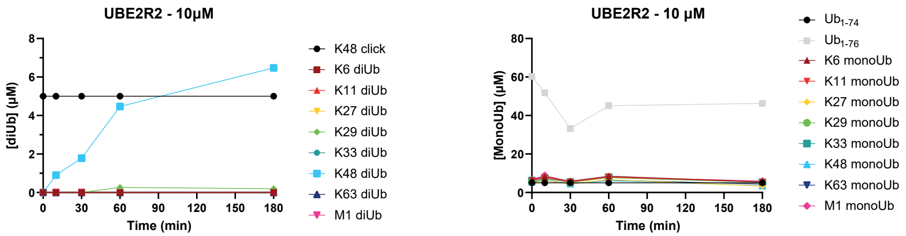


#### *UBE2G1*

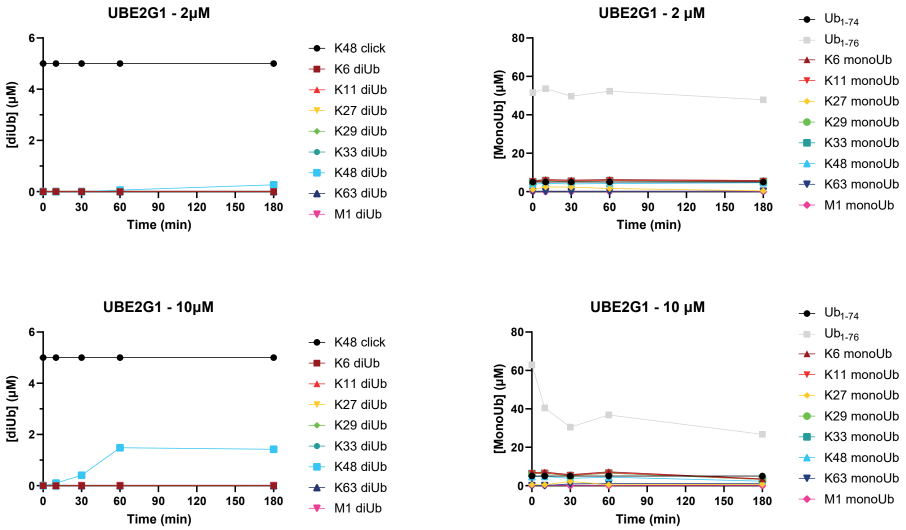


#### *UBE2N + UBE2V1*

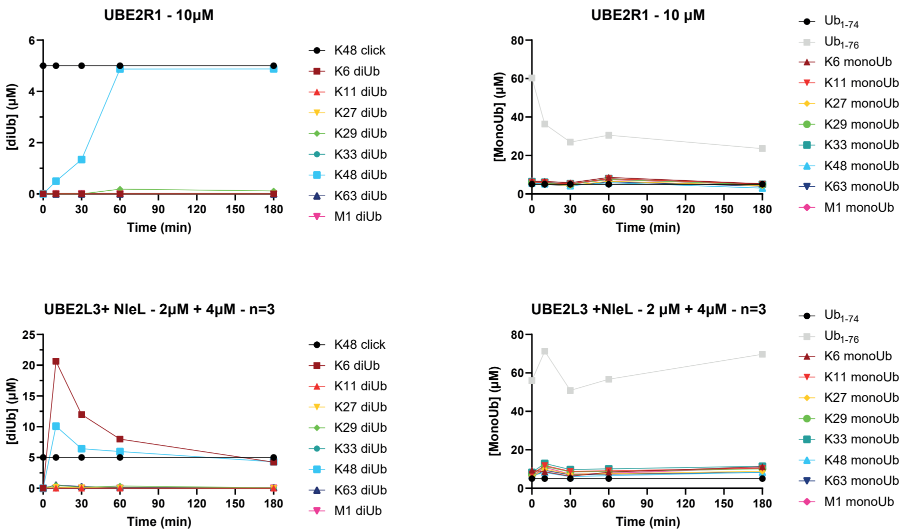

#### *UBE2D3 + HOIP*

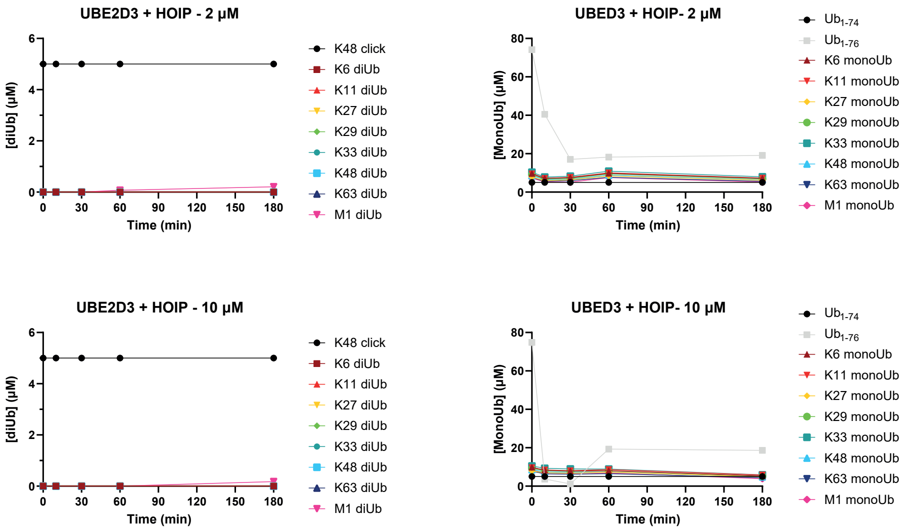


**Supplementary Figure 4| Determination of the assembly of all different diUb linkages built by specific E2 conjugating enzymes and E2-E3 pairs.** The quantified assay results are plotted. The first graph shows the amount of diUb (µM) formed in the reaction mixture normalized to the internal standard non-hydrolysable Lys48-linked diUb and calculated using the concentration of the internal standard as reference. The second graph shows the amount of monoUb (µM) present in the reaction mixture, normalized to the internal standard $Ub_{1-74}$ and calculated using the concentration of the internal standard as reference.

### *UBE2N + UBE2V1 + RNF4*

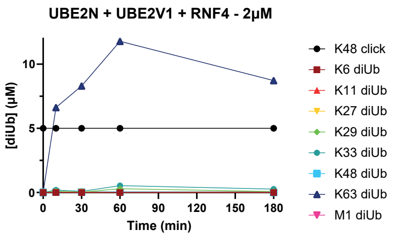


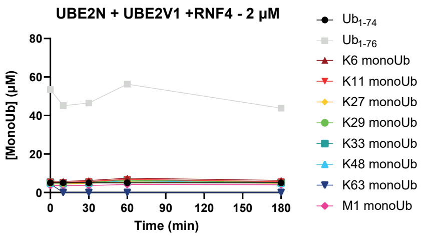


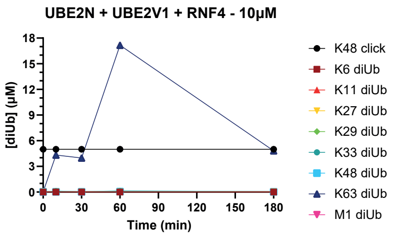


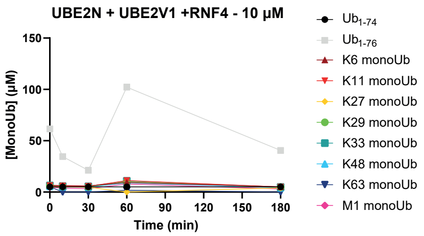


### *UBE2N + UBE2V1 + TRIM25*

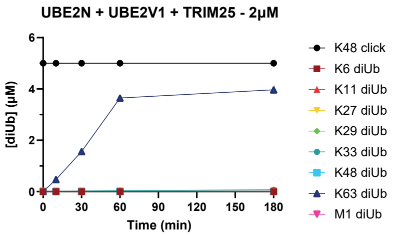


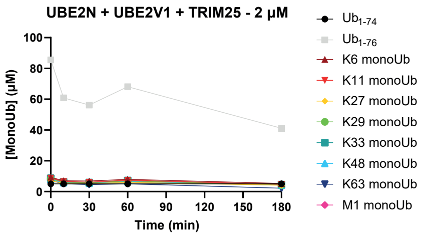


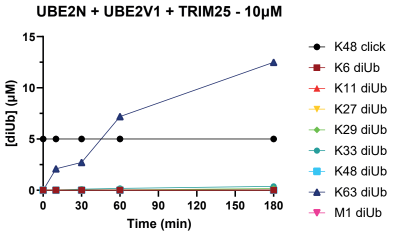


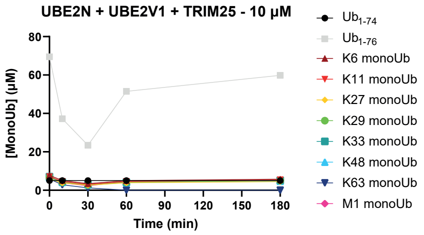


### *UBE2D3 + TRIM25*

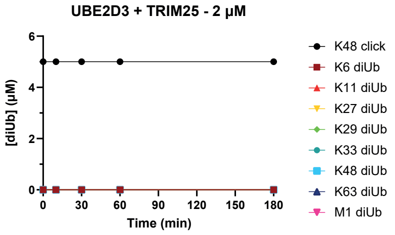


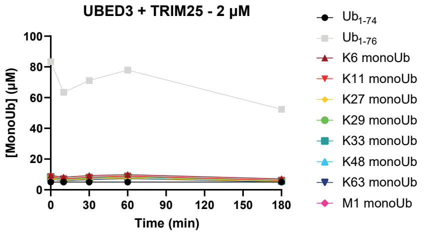

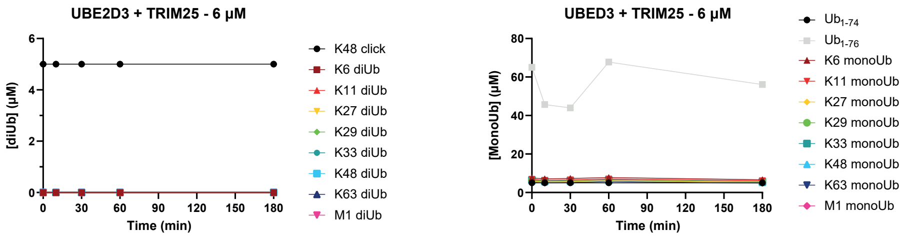


### UBE2D1 + RNF4

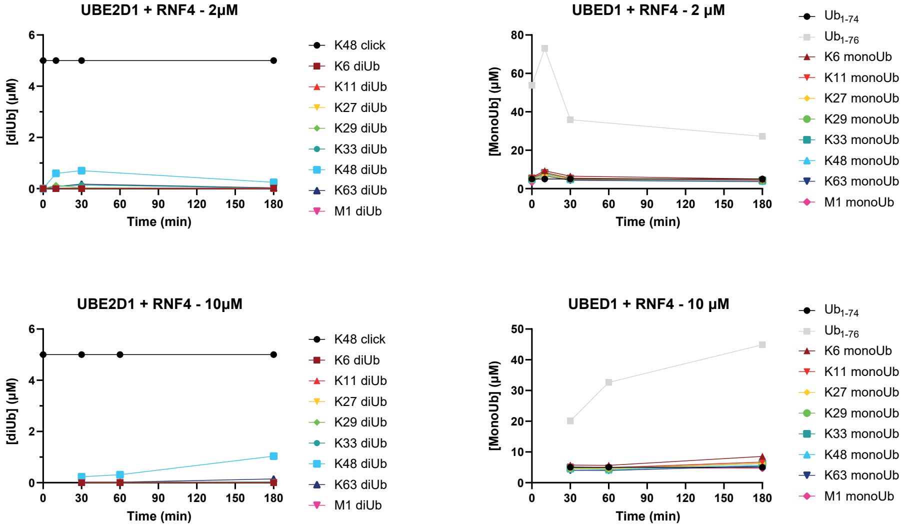


### UBE2D2 + RNF4

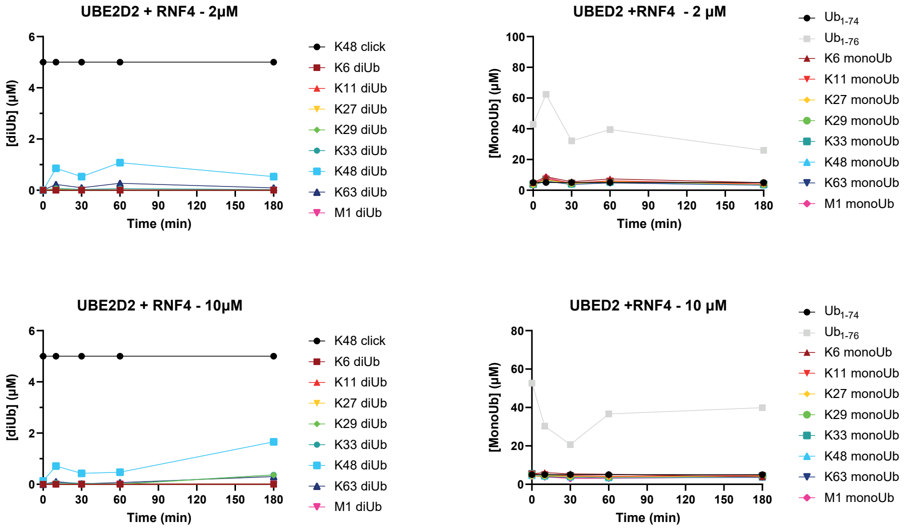

### *UBE2D3 + RNF4*

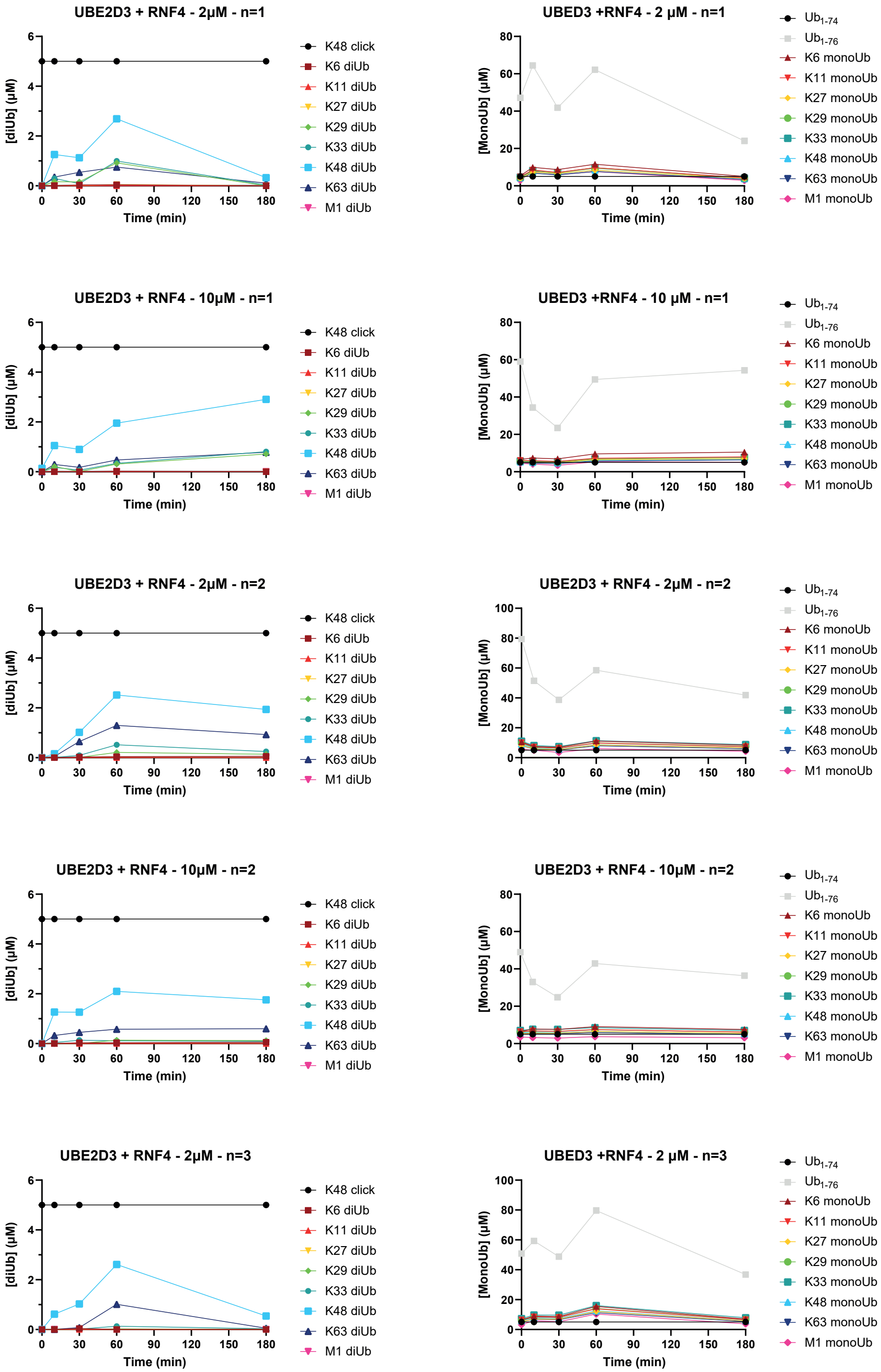
UBE2D3 + RNF4 - 2µM - n=1
[diUb] (µM)
Time (min)
K48 click
K6 diUb
K11 diUb
K27 diUb
K29 diUb
K33 diUb
K48 diUb
K63 diUb
M1 diUb
UBED3 +RNF4 - 2 µM - n=1
[MonoUb] (µM)
Time (min)
$Ub_{1-74}$
$Ub_{1-76}$
K6 monoUb
K11 monoUb
K27 monoUb
K29 monoUb
K33 monoUb
K48 monoUb
K63 monoUb
M1 monoUb
UBE2D3 + RNF4 - 10µM - n=1
UBED3 +RNF4 - 10 µM - n=1
UBE2D3 + RNF4 - 2µM - n=2
UBE2D3 + RNF4 - 2µM - n=2
UBE2D3 + RNF4 - 10µM - n=2
UBE2D3 + RNF4 - 10µM - n=2
UBE2D3 + RNF4 - 2µM - n=3
UBED3 +RNF4 - 2 µM - n=3

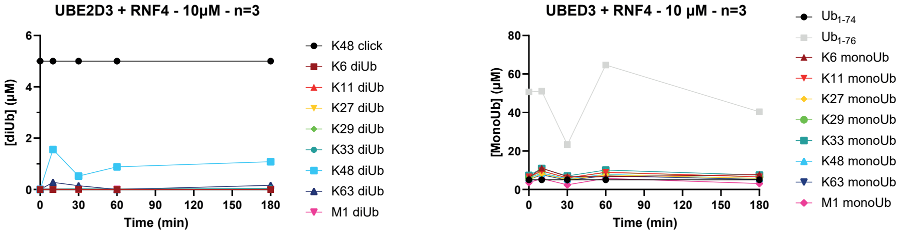


**Supplementary Figure 5| Determination of the assembly of diUb linkages built by specific E2 conjugating enzymes combined with different RING ligases.** The quantified assay results are plotted. The first graph shows the amount of diUb (μM) formed in the reaction mixture normalized to the internal standard non-hydrolysable Lys48-linked diUb and calculated using the concentration of the internal standard as reference. The second graph shows the amount of monoUb (μM) present in the reaction mixture, normalized to the internal standard $Ub_{1\text{-}74}$ and calculated using the concentration of the internal standard as reference.
